# Regulation of human interferon signaling by transposon exonization

**DOI:** 10.1101/2023.09.11.557241

**Authors:** Giulia Irene Maria Pasquesi, Holly Allen, Atma Ivancevic, Arturo Barbachano-Guerrero, Olivia Joyner, Kejun Guo, David M. Simpson, Keala Gapin, Isabella Horton, Lily Nguyen, Qing Yang, Cody J. Warren, Liliana D. Florea, Benjamin G. Bitler, Mario L. Santiago, Sara L. Sawyer, Edward B. Chuong

**Affiliations:** BioFrontiers Institute and Department of Molecular, Cellular & Developmental Biology, University of Colorado Boulder, Boulder, CO, 80309; Crnic Institute Boulder Branch, BioFrontiers Institute, University of Colorado Boulder, Boulder, CO, 80303; Division of Infectious Diseases, Department of Medicine, University of Colorado Anschutz Medical Campus, Aurora, CO, 80045; Division of Reproductive Sciences, Department of Obstetrics and Gynecology, University of Colorado Anschutz Medical Campus, Aurora, CO, 80045; Fred Hutchinson Cancer Research Center, Seattle, WA, 98109; The Ohio State University College of Veterinary Medicine, Columbus, OH, 43210; McKusick-Nathans Institute of Genetic Medicine, Johns Hopkins University School of Medicine, Baltimore, MD, 21205

**Keywords:** type I IFN, innate immunity, transposable elements, alternative splicing

## Abstract

Innate immune signaling is essential for clearing pathogens and damaged cells, and must be tightly regulated to avoid excessive inflammation or autoimmunity. Here, we found that the alternative splicing of exons derived from transposable elements is a key mechanism controlling immune signaling in human cells. By analyzing long-read transcriptome datasets, we identified numerous transposon exonization events predicted to generate functional protein variants of immune genes, including the type I interferon receptor IFNAR2. We demonstrated that the transposon-derived isoform of IFNAR2 is more highly expressed than the canonical isoform in almost all tissues, and functions as a decoy receptor that potently inhibits interferon signaling including in cells infected with SARS-CoV-2. Our findings uncover a primate-specific axis controlling interferon signaling and show how a transposon exonization event can be co-opted for immune regulation.

## INTRODUCTION

Innate immune signaling pathways orchestrate cellular responses to infection and diseases including cancer^1^. Impaired signaling underlies weakened responses to infection, while overactive responses lead to inflammatory diseases and many autoimmune disorders^2–5^. Consequently, these pathways are under tight regulatory control, yet we do not have a comprehensive understanding of these regulatory mechanisms. Although the primary immune signaling pathways are conserved in mammals, their response dynamics exhibit significant variation within and across species, implying that species-specific mechanisms underlie differences in immune regulation and function^6–8^. Although our understanding of these mechanisms is still in its infancy, elucidating the molecular basis of immune regulatory variation is key to understanding and treating immune-mediated diseases.

Transposable elements (TEs) are genetic “parasites” that constitute over 50% of the human genome^9^. Recently, we and others have shown that TEs are a source of cis-regulatory elements that control immune-inducible expression of immune genes, supporting a role for non-coding TEs in immune regulatory evolution^10–16^. Here, we investigated the impact of TE-derived exons on the protein-coding transcriptome, focusing on immune genes. Many intronic TEs contain splicing signal sequences that drive their incorporation into cellular transcripts, a process known as TE exonization^17^. Indeed, thousands of TE exonization events have been cataloged in the human genome based on transcriptomic data^18–21^. However, despite their potential to drive functional isoform diversification, most exonized TEs are silenced or weakly spliced, and assumed to be untranslated and nonfunctional^18,22,23^.

The advent of long-read transcriptomic technologies provides an unprecedented opportunity to discover and characterize full-length isoforms, including those derived from TE exonization events^24,25^. Here, we analyzed long-read transcriptomic datasets from human cells and uncovered hundreds of exonized TEs that generate alternative isoforms of immune genes that are robustly expressed. This included a truncated isoform of the Interferon Alpha and Beta Receptor subunit 2 (*IFNAR2*) generated by an exonized primate-specific Alu element. We functionally characterized this truncated isoform in human cells and found that it functions as a potent decoy receptor of IFN signaling, inhibiting IFN-mediated cytotoxicity and antiviral responses to SARS-CoV-2 and dengue virus. Our findings show how TE exonization generates new isoforms that can facilitate the regulation and evolution of immune signaling.

## RESULTS

### Multiple immune genes express TE-derived transcript isoforms

To explore the impact of TE exonization on the evolution of transcript isoforms of immune genes, we analyzed long-read transcriptomic datasets published for macrophages^26^ and healthy human tissues^27^ (Table S1). We used long-read transcriptome assemblies to screen for isoforms of protein-coding genes that contained at least one exonized TE and exhibited strong evidence of expression in at least one sample (Fig. 1A; for details, refer to the Methods section). This analysis revealed 3503 exonization events in 5991 isoforms for 2707 distinct genes (Table S1). Our analysis identified SINE transposable elements –Alu in particular– as the main class of transposable elements to undergo exonization both when inserted in the same (sense) or opposite (antisense) orientation of the protein coding gene they provide the alternative splice site to, followed by LINEs and LTR or DNA elements (Fig. 1B and Fig. S1). We then focused our analysis on genes involved in immune and inflammatory responses, which included 470 TE exonization events affecting 893 isoforms within 355 distinct genes. Many of these isoforms were predicted to encode truncated or altered protein variants that we reasoned could potentially functionally interfere with canonical protein activity. Additionally, nearly all these isoforms were primate-specific and either novel or poorly characterized.

**Fig. 1.**
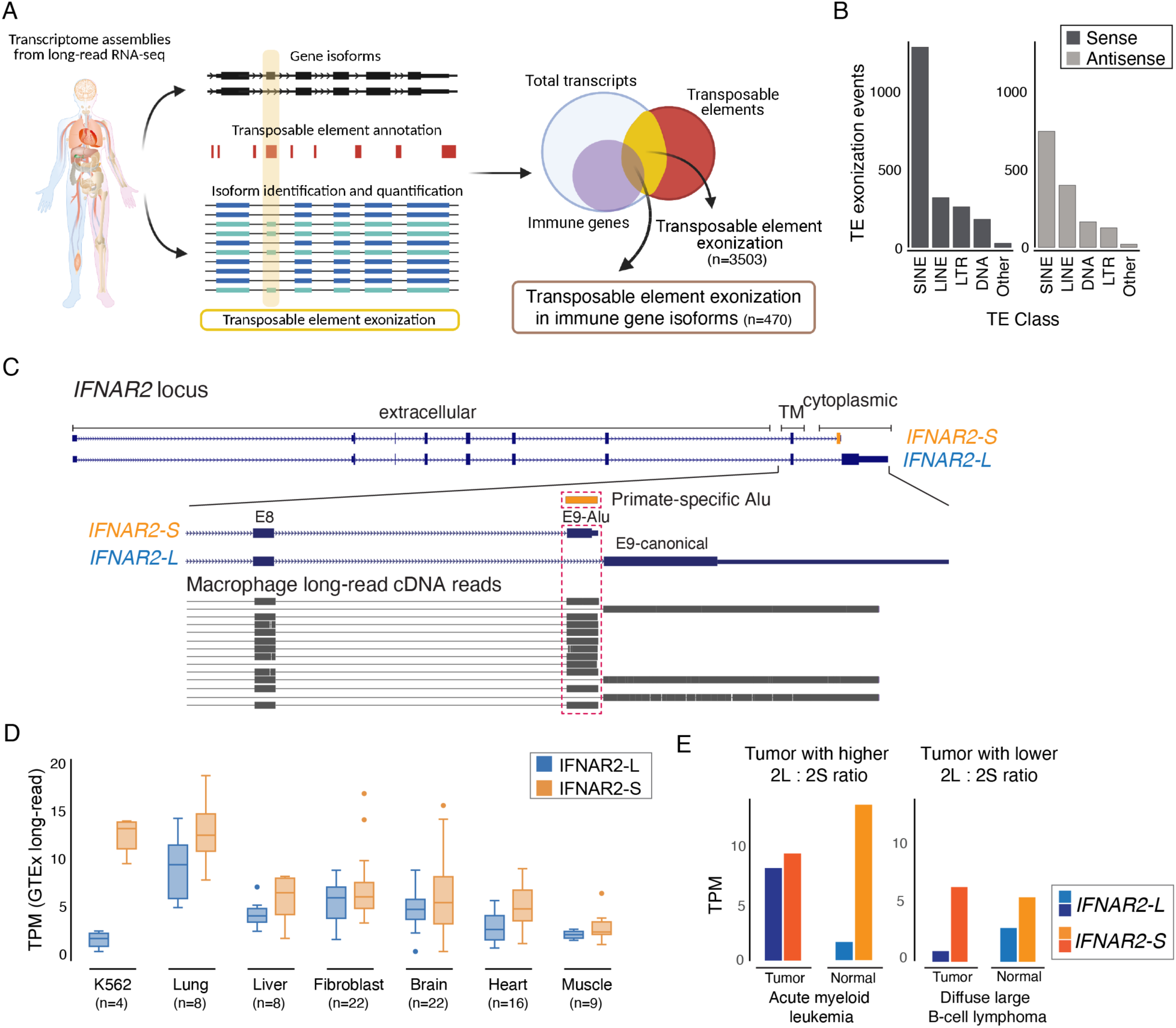
Human IFNAR2 is constitutively spliced into two main isoforms. **A)** Schematic of the approach to identify candidate exonized transposable elements (TEs) in immune genes from long-read RNA sequencing data. **B)** Count of TE exonization events for each main TE class based on their orientation compared to that of the protein-coding gene they provide the alternative splice site to (sense= same orientation, to the left; antisense = opposite orientation, to the right). Only TEs that overlapped a protein-coding gene exon for 80% of their length and a splice site for isoforms with TPM > 5 in at least one sample were included^27^. **C)** Read alignments from long-read cDNA sequencing data of human macrophages^26^ over the short IFNAR2 alternative isoform (*IFNAR2-S*) and the canonical long IFNAR2 isoform (*IFNAR2-L*). **D)** Long-read RNA-seq read counts comparing *IFNAR2-S* (orange) and *IFNAR2-L* (blue) expression across normal human tissues (GTEx^27^). **E)** Expression levels of *IFNAR2-S* and *IFNAR2-L* in matched healthy (lighter shade) and tumor (darker shade) samples in acute myeloid leukemia and diffuse large B-cell lymphoma (TCGA^33,90^). The two tumor types show extreme changes in the relative isoform expression levels compared to matched healthy samples: increase in *IFNAR2-L* relative expression in leukemia, and decrease in *IFNAR2-L* relative expression in lymphoma. TM = transmembrane domain. TPM = Transcripts per million.

We focused on a truncated short isoform of the receptor IFNAR2 (NM_000874, *IFNAR2-S* hereafter), formed by the exonization of a primate-specific Alu element within the last intron of the gene (Fig. 1C and Fig. S2). The canonical full-length isoform of IFNAR2 (NM_001289125, *IFNAR2-L* hereafter) encodes one of the core receptors for type I interferons (IFNs), which are cytokines secreted by infected or damaged cells. Upon binding to IFN, IFNAR2-L forms a complex including IFNAR1, which transduces intracellular activation of STAT1/2-dependent transcription of immune and inflammatory genes^28^.

*IFNAR2-S* is predicted to generate a truncated receptor that lacks the intracellular signaling domain^29^, and we speculated that it could act as a decoy receptor that sequesters the ligand and inhibits signaling. *IFNAR2-S* was previously identified thirty years ago during the initial cloning of human IFNAR2^30^, but remains functionally uncharacterized except for one early study using transfected mouse cells^31^, which lack this isoform. To date, research on human IFN signaling has exclusively focused on the canonical *IFNAR2-L* isoform. Given that our long-read analysis supported robust expression of the alternative *IFNAR2-S* isoform, we sought to investigate the physiological significance of IFNAR2-S in human cells.

### IFNAR2-S is the predominantly expressed isoform of IFNAR2 in human cells

To determine the tissue-specificity of IFNAR2-S expression, we quantified isoform expression across a panel of long-read and short-read RNA-seq datasets for healthy^27,32^ and cancer tissues^33^. Our analysis revealed that both *IFNAR2-L* and *IFNAR2-S* isoforms are constitutively expressed in nearly all tissues examined. Surprisingly, we found that *IFNAR2-S* is expressed at equivalent or higher levels of *IFNAR2-L* across healthy tissues, based on both full-transcript quantification from long-read RNA-seq data (Fig. 1C-D) and junction read counts from short-read RNA-seq data (Fig. S3A). Given that the canonical *IFNAR2-L* isoform is widely assumed to be the main transcript expressed in human cells, our finding suggests that *IFNAR2-S* may play a role in human IFN signaling.

We also examined isoform expression in short-read RNA-seq data from matched normal and cancer samples profiled by the Cancer Genome Atlas (TCGA^33^; Fig. S3B). We observed expression of both *IFNAR2-S* and *IFNAR2-L* in all samples, with notable shifts in the relative isoform ratio (*IFNAR2-L* : *IFNAR2-S*) when comparing tumor and normal samples for several cancer types, including diffuse Large B-cell lymphoma and Acute Myeloid Leukemia (Fig. 1E). Therefore, although both isoforms share the same transcription start site, the relative expression levels of *IFNAR2-L* and *IFNAR2-S* are subject to dysregulation in disease states.

### IFNAR2-S encodes a nonsignaling type I IFN receptor

Our transcriptomic analyses were consistent with a potential functional role for *IFNAR-S* in immune regulation. We investigated this possibility using human cervical adenocarcinoma HeLa cells, which are a widely used model for studying type I IFN signaling^34–38^, and human pulmonary carcinoma A549 cells, which are widely used to study the role of IFN in the context of infections^39^. Similar to most human tissues, HeLa cells show higher expression of *IFNAR2-S* than *IFNAR2-L* (Fig. S4B), and do not constitutively express type I IFNs. We confirmed that peptides unique to IFNAR2-S were detected in mass spectrometry data (Fig. S5), indicating that *IFNAR2-S* is translated. We also experimentally verified that the endogenous *IFNAR2-S* isoform is translated into a stable protein by using CRISPR to tag the Alu exon with a small HiBiT epitope tag, and detected the tagged protein at the expected size (Fig. S6; see Supplementary materials and methods).

To disentangle the functions of the short and long IFNAR2 isoforms, we used CRISPR to generate multiple isoform-specific knockout (KO) clones by deleting either canonical exon 9 (E9-canonical) to generate the IFNAR2-L KO or the Alu-derived terminal exon (E9-Alu) to generate the IFNAR2- S KO. We also generated a full IFNAR2 KO by deleting the shared exon 7 (E7) which introduces an early frameshift (Fig. 2A and Fig. S4; see Supplementary materials and methods). To orthogonally validate results from our isoform-specific KO clones, we designed isoform-specific short interfering RNAs (siRNAs) targeting E9-canonical or E9-Alu (Fig. S4D-F).

**Fig. 2.**
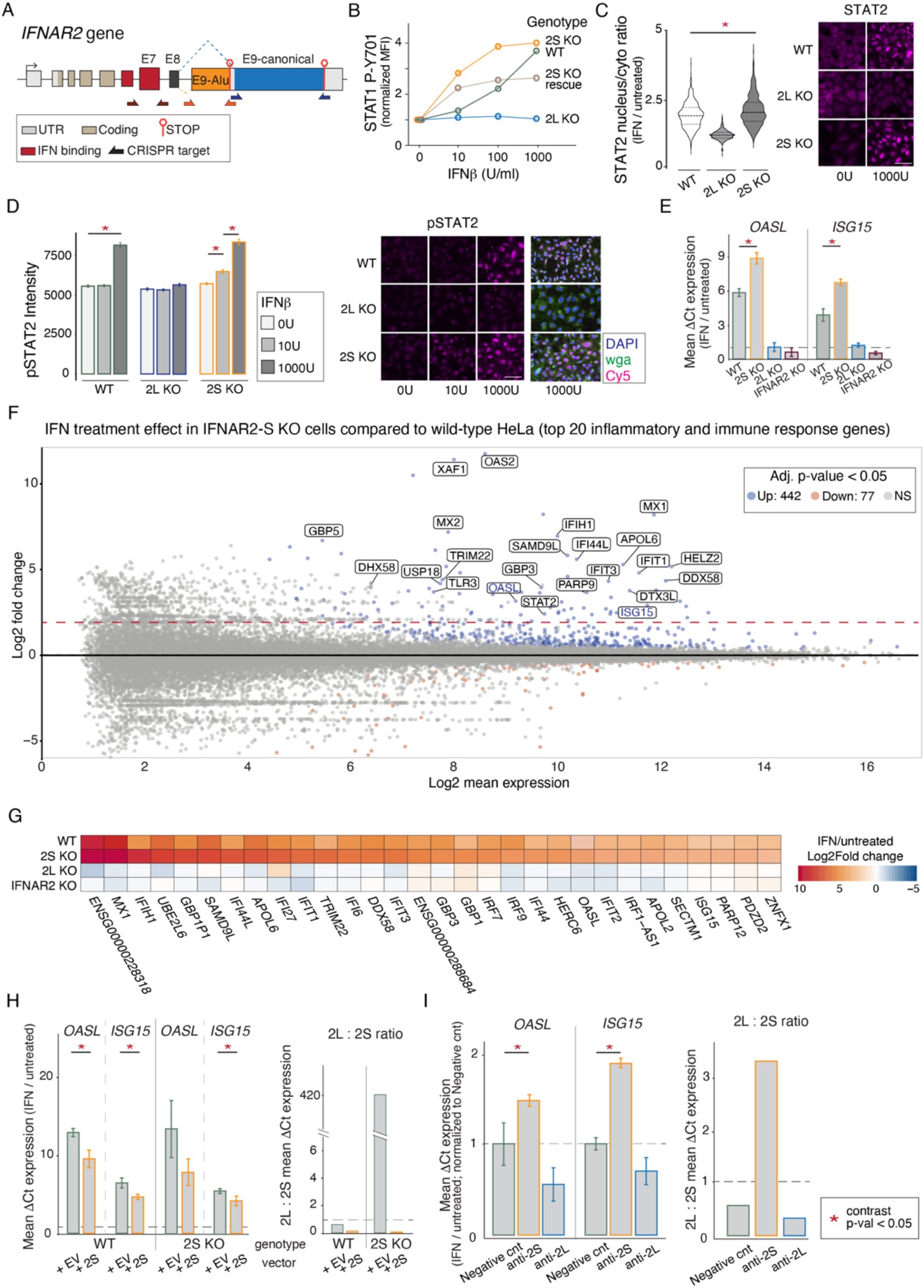
IFNAR2-S functions as a decoy receptor for type I IFN signaling. **A)** Schematic model of isoform-specific knockouts (KOs) of the *IFNAR2* gene (not drawn to scale). IFNAR2-S KO cell lines were generated by deleting the E9-Alu exon (orange); IFNAR2-L KO cell lines were generated by deleting the E9-canonical exon (blue); complete IFNAR2 KO cell lines were generated by deleting exon 7 (E7 KO/IFNAR2 KO). **B)** Plot shows changes in normalized STAT1 phosphorylation levels as detected by phospho-flow cytometry in wild-type and knockout HeLa cell lines upon 30min treatment with increasing doses of IFNβ. Data shown are mean fluorescence intensity (MFI) normalized against untreated cells of the same genotype. **C-D)** STAT2 and phosphorylated STAT2 (pSTAT2) signal from wild-type and knockout A549 cell line. On the left, **C**: violin plots show the ratio of nuclear to cytoplasmic STAT2 signal in cells treated with 1000U/ml of IFNβ normalized to untreated cells (wild-type n=663, IFNAR2-L KO n=377, IFNAR2-S n=574); **D**: histograms show total pSTAT2 signal in untreated cells and cells treated with 10U/ml and 1000U/ml of IFNβ. On the right, immunofluorescence pictures of STAT2 and pSTAT2 from which fluorescence signals were quantified. Pink = (p)STAT2 signal. Scale bar= 50μm. **E)** RT-qPCR measurements of *OASL* and *ISG15* fold-change induction comparing untreated to 4hrs 10U/ml IFNβ treatment in HeLa wild-type and KO cell lines. For each genotype, 3 biological replicates (different KO clonal cell lines) were tested and assayed in triplicate. **F)** RNA-Seq MA-plot showing differences in 4hrs 10U/ml IFNβ treatment comparing wild-type and IFNAR2-S KO HeLa cells. Significantly differentially induced genes with annotated immune or inflammatory functions (GO:0006955 and GO:0006954) are labeled. **G)** Heatmap of the top 30 most significantly upregulated genes (based on adjusted p-value) by IFNβ in IFNAR2-S KO cells. Displayed are the log2fold change values from treated vs untreated pairwise comparisons for each HeLa cell line. **H)** RT-qPCR measurements of *OASL* and *ISG15* fold-change induction comparing untreated to 4hrs 10U/ml IFNβ treatment in wild-type or IFNAR2-S KO cells overexpressing empty vector or IFNAR2- S. **I)** RT-qPCR measurements of *OASL* and *ISG15* fold-change induction in response to a 4hrs 10U/ml IFNβ treatment in wild-type HeLa cells transfected with negative control, anti-IFNAR2-L, or anti-IFNAR2-S siRNAs. Asterisks (*) indicate a significant difference in the effect of treatment (p-value < 0.05) between conditions as calculated through an ANOVA *emmean* pairwise contrast statistical test.

First, we tested how each IFNAR2 isoform affects JAK/STAT signaling in response to IFN stimulation. IFNAR2-S lacks the canonical intracellular domain responsible for phosphorylation of STAT2 and STAT1^28^. Accordingly, we confirmed that IFNAR2-L, but not IFNAR2-S, was required for phosphorylation of STAT1/2 upon type I IFN treatment (Fig. S7A). We next quantified phosphorylated STAT1 (pSTAT1) levels as a function of IFNβ concentration by phospho-flow cytometry. Wild-type cells showed an expected increase in STAT1 phosphorylation in response to IFNβ (up to 2000U/ml), while IFNAR2-L KO cells showed no response to the treatment. Notably, IFNAR2-S KO cells showed an amplified response compared to wild-type cells at lower IFN concentrations, with roughly 2-fold greater induction of pSTAT1 at 10 and 100U/ml IFNβ and response saturation at 100U/ml (Fig. 2B and Fig. S7B). Overexpression of IFNAR2-S in IFNAR2- S KO cells partially rescued IFN response dynamics to those observed in wild-type cells.

We confirmed these results by assessing STAT2 phosphorylation and nuclear translocation by immunofluorescence (IF) in wild-type and knockout A549 cell lines. Upon treatment with IFNβ, we observed significantly higher nuclear:cytoplasm STAT2 fluorescence signal in IFNAR2-S KO cells compared to wild-type cells (2-way ANOVA p-value < 0.001), whereas no significant changes in STAT2 localization were observed in IFNAR2-L KO cells (Fig. 2C). We also measured pSTAT2 fluorescence signal at different IFNβ concentrations (Fig. 2D and Fig. S8). Similarly to what observed in HeLa cells, we detected significantly higher STAT2 phosphorylation in IFNAR2-S KO compared to wild-type A549 cells even at lower doses of IFNβ treatment (10U/ml). Together, these results demonstrate that the presence of IFNAR2-S strongly inhibits JAK/STAT activation in response to sub-saturating levels of type I IFN.

### IFNAR2-S inhibits ISG activation

We next tested whether inhibition of JAK/STAT signaling by IFNAR2-S affects downstream activation of interferon stimulated genes (ISGs). We treated wild-type and KO cell lines with 10U/ml of IFNβ or a mock treatment for 4hrs, and assayed fold-change induction of the canonical ISGs *OASL* and *ISG15* by RT-qPCR. As expected, we observed robust ISG induction in wild-type cells but no ISG induction in IFNAR2-L KO cells and complete IFNAR2 KO cells. Compared to wild-type cells, IFNAR2-S KO cells showed significantly higher fold-induction of ISGs upon IFN stimulation (Fig. 2E; p-value *OASL* < 0.0001, p-value *ISG15* = 0.022; Table S2). Overexpression of IFNAR2-S in either wild-type or IFNAR2-S KO cells was sufficient to dampen ISG induction (Fig. 2H; Table S2). We replicated these experiments in wild-type cells where each isoform was transiently silenced using exon-specific siRNAs, and confirmed that silencing IFNAR2-S led to increased ISG induction (Fig. 2I; Table S2). These experiments indicate that IFNAR2-S inhibits IFN-mediated transcriptional activation of ISGs.

To extend our analysis to a genome-wide level, we used RNA-seq to profile ISG activation in wild-type, IFNAR2-L KO, IFNAR2-S KO and complete IFNAR2 KO cells (see Supplementary materials and methods). In wild-type cells, we identified 94 ISGs (adj. p-value < 0.05, log2fold change > 2) (Fig. S9; Table S3). We confirmed that IFNAR2-L KO and full IFNAR2 KO cells failed to respond to IFN stimulation (Fig. S10; Table S3). Consistent with our RT-qPCR results, IFNAR2-S KO cells showed a global increase in ISG activation (Fig. S11; Table S3). We tested for the effect of IFN treatment between IFNAR2-S KO and wild-type cells, and identified 85 genes that showed significantly higher induction in IFNAR2-S KO cells (Fig. 2F; adj. p-value < 0.05, log2fold change > 2; Table S3). These predominantly included genes associated with the response to type I IFN (i.e., IFI family genes, IRF family genes, IFIT family genes, *OAS1*, *STAT2* and *MX* genes), response to viruses (i.e., *PARP9*, *TRIM22, TRIM34* and *TLR3*), binding to double-stranded RNAs (i.e., *DDX58* and *DHX58*), RIG-I-like and NOD-like receptor signaling pathways, and apoptosis (i.e., *TNFSF10*). Only 23 genes – mostly non-coding RNAs – were more highly induced in wild-type cells, which may be due to off-target differences between clones (Table S3). Taken together, these results demonstrate that IFNAR2-S functions to restrain the transcriptional activation of ISGs, and the absence of IFNAR2-S leads to genome-wide increase in the fold-change of ISG induction following IFNβ stimulation (Fig. 2G).

### IFNAR2-S encodes a decoy receptor for the type I IFN response

We next investigated how IFNAR2-S affects binding and internalization of IFN using HeLa cells, which do not produce IFNβ. In cells treated with 1000U/ml of IFNβ for 30min, we detected IIFNβ in the lysates of wild-type, IFNAR2-S KO and IFNAR2-L KO cells, but not in full IFNAR2 KO cells (Fig. 3A), indicating that both IFNAR2-L and IFNAR2-S are able to bind type I IFN. We further quantified IFNβ levels in HeLa cell lysates by ELISA, and found that upon IFNβ treatment IFNAR2- S KO cells exhibited the highest levels of internalized IFN, followed by wild-type and IFNAR2-L KO cells, with the lowest levels in full IFNAR2 KO cells (Fig. S12A). We also measured the residual levels of IFNβ in the media, and confirmed that IFNAR2-S KO cells had the lowest levels of soluble IFNβ, consistent with increased uptake driven by the expression of IFNAR2-L alone (Fig. S12B). Together, these results indicate that while both isoforms can bind IFN, there may be differences in the receptor and/or cytokine trafficking based on domains specific to the different intracellular regions^40^.

**Fig. 3.**
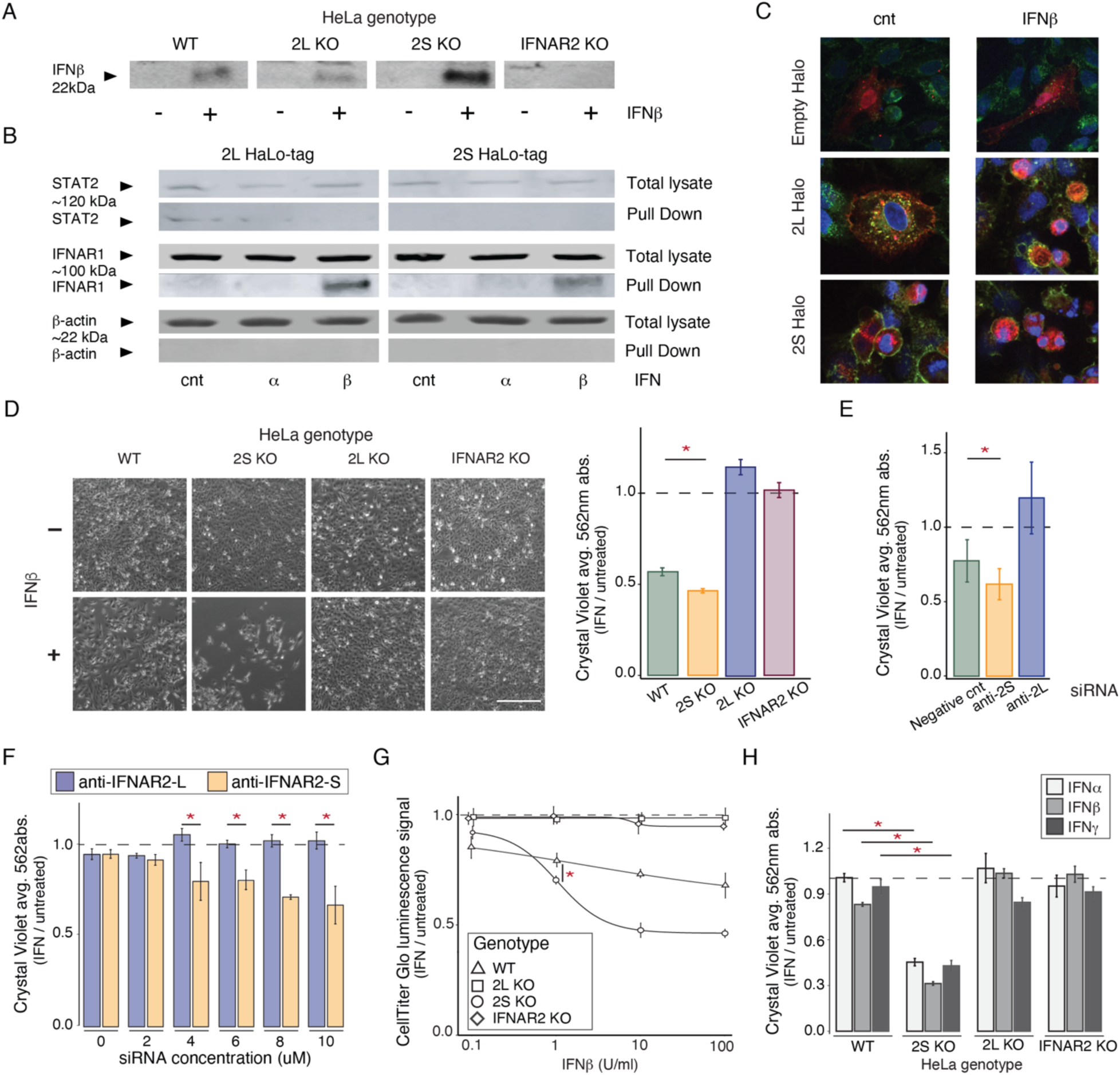
IFNAR2-S acts as a decoy receptor for the type I IFN signaling. **A)** Western blot of IFNβ in lysates of wild-type and knockout HeLa cells either untreated or treated with 1000U/ml IFNβ for 30min shows that both IFNAR2 isoform can bind IFNβ. **B)** Western blots of STAT2, IFNAR1 and β-actin in total lysates and pull-down fractions of IFNAR2 KO HeLa cells overexpressing either the IFNAR2-S or the IFNAR2-L isoform tagged with a C-terminus HaloTag. The HaloTag was used to perform isoform-specific pull-downs to identify interactors of IFNAR2-L and IFNAR2-S independently in untreated cells and cells treated with IFNα for 10min or IFNβ for 5min. Target proteins were detected in total cell lysates (pre-pull-down fraction) in all treatment conditions. STAT2 was detected only in IFNAR2-L HaloTag pull-downs, and IFNAR1 in cells treated with IFNβ. Signal was barely detectable in cells treated with IFNα, consistent with the lower affinity of IFNα to IFNAR2. β-actin was not identified in the pull-down fractions. **C)** Immunofluorescent labeling of IFNAR2-L-HaloTag and IFNAR2-S-HaloTag in untreated conditions (cnt, on the left) and after 30min of IFNβ treatment (100U/ml). Compared to the diffused signal of cells expressing an empty HaloTag vector, both cell lines expressing IFNAR2-L-HaloTag and IFNAR2-S-HaloTag show clear punctuated signal in the cytoplasm and at the cell membrane. Magnification = 40x; Green = wga; membrane staining; Blue = DAPI, nucleus; Red = permeable JFX549 HaloTag ligand. **D)** Brightfield images of HeLa cells at 4x magnification either untreated or treated with 10U/ml IFNβ for 4 days (left) and matched normalized cell viability measured by crystal violet staining for the same cells. Scale bar = 1000μm. **E)** Normalized cell viability measured by crystal violet staining of wild-type cells transfected with 5μM negative control, anti-IFNAR2-L, and anti-IFNAR2-S siRNAs and treated with 10U/ml IFNβ for 4 days. **F)** Normalized cell viability measured by crystal violet staining of wild-type cells transfected with increasing concentration of scrambled, anti-IFNAR2-L, and anti-IFNAR2-S siRNAs and treated with 10U/ml IFNβ for 4 days. **G)** Dose-response curves of normalized cell viability measured by CellTiter-Glo in response to treatment with increasing doses of IFNβ (0 to 100U/ml). **H)** Cell viability measured by crystal violet staining of wild-type and KO cells treated with 100U/ml of IFNα, IFNβ and IFNγ for 7 days. All treatments were performed in triplicate and depicted as mean and standard error and normalized to untreated controls. Asterisks (*) indicate a significant difference in the effect of treatment (p-value < 0.05) between conditions as calculated through an ANOVA *emmean* pairwise contrast statistical test.

To determine how IFNAR2-S interferes with the signaling complex formation, we overexpressed either IFNAR2-S or IFNAR2-L fused with a C-terminal HaloTag in wild-type and KO cells to determine interactions with IFNAR1 and STAT2 (Fig. S13). We first verified that addition of the HaloTag does not disrupt IFNAR2-L activation of JAK/STAT signaling by IFNβ (Fig. S14). We then pulled down IFNAR2-L-HaloTag or IFNAR2-S-HaloTag and their associated proteins in cells either untreated or treated with IFNβ. We detected IFNAR1 in pull-down fractions of both IFNAR2- S-HaloTag and IFNAR2-L-HaloTag in IFN-treated cells, confirming that IFNAR2-S can also form a ternary complex with IFNAR1 and IFNβ. Moreover, while we detected STAT2 in the IFNAR2-L-HaloTag pull-down from untreated cells as expected^36,38^, we did not detect STAT2 association with IFNAR2-S-HaloTag (Fig. 3B). These results indicate that IFNAR2-S is a decoy receptor that forms a nonproductive ternary complex with IFN and IFNAR1 that inhibits cellular activation of JAK/STAT signaling.

Finally, we used fluorescent ligands to label the IFNAR2-L-HaloTag and IFNAR2-S-HaloTag fusion proteins and visualized them by confocal imaging. In IFNβ-stimulated cells, IFNAR2-L was present in cytoplasmic puncta that co-localize with Rab5, a key regulator of endosome fusion and trafficking_40_. IFNAR2-S-HaloTag showed similar localization patterns, consistent with the retention of the transmembrane domain (Fig. 3C and Fig. S13). Combined, these results suggest that IFNAR2-S acts as a decoy receptor that directly competes with IFNAR2-L for type I IFN binding and association with IFNAR1.

### IFNAR2-S restrains type I IFN-induced cytotoxicity

Having established that IFNAR2-S acts as a decoy receptor that inhibits JAK/STAT signaling and ISG activation by type I IFN, we next investigated the functional consequences of IFNAR2-S. We assayed IFNβ-induced cytotoxicity^34^ by measuring cell proliferation and viability after prolonged IFN exposure (10U/ml for 4 days). Both wild-type and IFNAR2-S KO HeLa cells showed significantly reduced cell viability in response to IFN treatment (Mann-Whitney U Test p-value <0.01 and 0.001, respectively; Fig. 3D; Table S4), but IFNAR2-S KO cells showed dramatically lower viability upon treatment (p-value = 0.002), consistent with our previous experiments. The increase in IFN-mediated cytotoxicity in IFNAR2-S KO cells was evident at a range of doses (0.1 to 100U/ml IFNβ; Table S4), and overexpressing IFNAR2-S in IFNAR2-S KO cells was sufficient to restore IFN sensitivity to near wild-type levels (Fig. S15). These results demonstrate that IFNAR2-S directly inhibits the cytotoxic effects of prolonged IFN exposure.

We observed concordant results in wild-type cells where isoforms were depleted using isoform-specific siRNAs prior to prolonged IFNβ exposure (Fig. 3E; Table S4). We modulated the relative ratio of *IFNAR2-L : IFNAR2-S* expression by transfecting different concentrations of isoform-specific siRNAs while keeping the total amount of transfected siRNA constant. We found that at higher siRNA doses (>4μM and <10μM) increasing the *IFNAR2-L : IFNAR2-S* ratio led to significantly lower cell viability (Fig. 3F; Table S4). Finally, overexpression of IFNAR2-S in both wild-type and IFNAR2-S KO cells was sufficient to reduce sensitivity to IFN-mediated cytotoxic effects (Fig. S15; Table S4). These results indicate that tuning the *IFNAR2-L : IFNAR2-S* ratio can modulate cellular responses to IFN.

We next tested how signaling by different interferons (IFNα, IFNβ, IFNγ) is impacted by the different IFNAR2 isoforms. We assayed cell viability for wild-type and KO HeLa cells after prolonged IFN exposure (100U/ml of each IFN for 7 days) (Fig. 3H, Fig. S16 and Tables S2 and S4). Wild-type cells showed reduced viability in response to IFNβ, but not IFNα (in contrast to Shi et al.^41^) or IFNγ. Interestingly, IFNAR2-S KO cells showed an increased loss of viability in response to the type I IFNs (IFNα and IFNβ) as expected, but also showed reduced viability in response to type II IFN (IFNγ) (Table S4), potentially due to signaling crosstalk between the type I and II IFN pathways^42^. These findings suggest that IFNAR2-S functions as a buffer against the cytotoxic effects of type I IFN as well as other signaling pathways that may indirectly activate IFN signaling.

We replicated our findings in additional human cell lines, including A549 lung adenocarcinoma cells and PEO-1 ovarian adenocarcinoma cells. In contrast to HeLa cells, A549 and PEO-1 cells express higher levels of *IFNAR2-L* compared to *IFNAR2-S* (∼2x, Fig. S4). Silencing IFNAR2-L in each cell line led to reduced ISG activation and increased viability in response to IFNβ, whereas silencing IFNAR2-S led to increased ISG activation and decreased cell viability (Fig. S17; Tables S2 and S4). Overexpression of IFNAR2-S in A549 cells also inhibited the cytotoxic effect of IFNβ in a dose-dependent manner (Fig. S17C; Tables S2 and S4). Together with our previous observation that IFNAR2-S is broadly expressed (Fig. 1D), our findings indicate that IFNAR2-S is a key regulator of IFN signaling that functions in virtually all human tissues.

### The IFNAR2-L : IFNAR2-S relative isoform ratio affects cell response to viral infection

Type I interferons are key contributors to the host antiviral response, therefore we investigated how IFNAR2-S affects immune responses to viral infection. First, we asked whether *IFNAR2-S* transcript expression changes during an immune response by analyzing RNA-seq time-course datasets of human cell lines responding to viral infection. In Calu3 cells infected by SARS-CoV-2_43_ we observed upregulation of *IFNAR2-L* (Fig. 4A, left) as soon as 3hrs post infection. Conversely, in A549 cells infected by Influenza B virus^44^, we detected downregulation of *IFNAR2- L* in response to infection (Fig. 4A, right). These findings indicate that the relative levels of the long and short *IFNAR2* isoforms are subject to distinct transcriptional dynamics upon viral infection, implying that isoform regulation of *IFNAR2* may contribute to modulating immune responses.

**Fig. 4.**
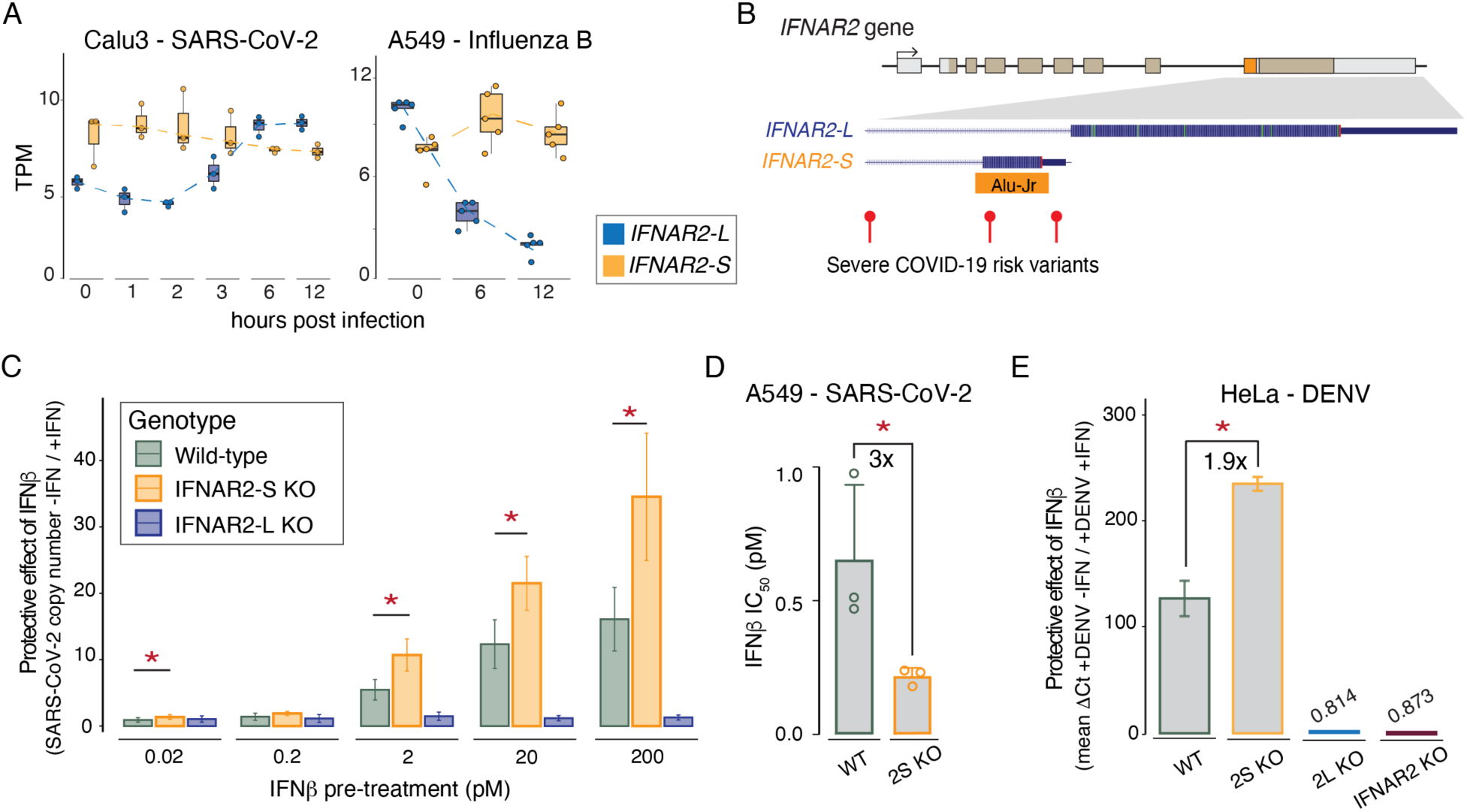
The IFNAR2-L : IFNAR2-S ratio affects cellular responses to viral infection. **A)** Box plots show dynamic modulation of *IFNAR2* isoform expression levels (TPM) in human cell lines infected with different viruses. SARS-CoV-2 data from (*42*) and Influenza B data from (*43*). **B)** Variants associated with severe COVID-19 located in and near exon E9-Alu as identified in GWAS studies by the COVID-19 Host Genetic Initiative^91^. **C)** Bar plot shows the protective effect of IFNβ in the context of SARS-CoV-2 (B.1.617.2) viral infection in wild-type and knockout A549-ACE2 cells. Each bar represents the ratio of viral genome copies in the media of cells that were not pre-treated with IFNβ over viral genome copies in the media of cells that were pre-treated with increasing doses of IFNβ. SARS-CoV-2 copy number was measured by RT-qPCR. **D)** Based on three independent SARS-CoV- 2 infection experiments, the IFNβ IC_50_ was calculated for wild-type and IFNAR2-S KO A549-ACE2 cells. Bars show the average and distribution of IC_50_ values for the two genotypes. **E)** Histogram shows the protective effect of IFNβ in the context of dengue virus 2 (DENV-2) viral infection in wild-type and knockout HeLa cells. Protective effect was calculated as the ratio of viral genome load detectable in lysates by RT-qPCR of unstimulated cells over the viral genome load in lysates of cells that were pre-stimulated with 100U/ml of IFNβ. Asterisks (*) indicate a significant difference in the effect of treatment (p-value < 0.05) between conditions as calculated through an ANOVA *emmean* pairwise contrast statistical test

We therefore investigated whether IFNAR2-S plays a role in the IFN-mediated antiviral response to SARS-CoV-2. Dysregulated type I IFN responses underlie many cases of severe COVID-19 ^45–47^. Genetic studies have linked over 75 single-nucleotide variants within the IFNAR2 locus to severe COVID-19^48,49^, but most have unknown significance and mechanism. We identified multiple variants located within the E9-Alu exon –but not within E9-canonical– that are associated with severe COVID-19 (Fig. 4B). These observations suggest that aberrant IFNAR2-S expression may be a causal mechanism underlying pathological responses to viral infection.

We generated IFNAR2-S KO and IFNAR2-L KO A549 cell lines that stably express the ACE2 receptor, which is required to support SARS-CoV-2 entry^39^ (Fig. S18A-C). We verified that A549- ACE2 IFNAR2-S KO cells show increased ISG activation upon IFNβ stimulation, consistent with results from HeLa KO cells (Fig. S18D). To assay the effect of IFN on viral replication, we primed wild-type and KO cells with varying doses of IFNβ prior to infection with SARS-CoV-2 virus (B.1.617.2), and compared viral RNA levels after 24hrs between primed and unprimed cells (Fig. S19, Table S5). As expected, IFN treatment had a clear protective effect by significantly reducing viral replication in wild-type cells. Strikingly, the protective effect of IFN was 2.5 times stronger in IFNAR2-S KO cells compared to wild-type cells at high doses (20-200pM, p-values = 0.005 and 0.002, respectively), and was also evident even at low doses (1.5x at 0.02pM, p-value = 0.033; Fig. 4C), leading to significantly reduced viral replication. IFNAR2-S KO cells also showed a significantly reduced IFNβ IC_50_ compared to wild-type cells (p-value = 0.035; Fig. 4D), indicating that in absence of IFNAR2-S lower doses of IFN can induce more potent antiviral responses to SARS-CoV-2. These experiments indicate that IFNAR2-S dampens the antiviral response to SARS-CoV-2, which may be important for preventing overactive IFN responses that contribute to severe COVID-19^45–47^.

We replicated our findings with another virus and cell line using dengue virus (DENV-2), which is restricted by type I IFN^50,51^ and can infect HeLa cells^52,53^. Using wild-type and KO HeLa cells, we infected unprimed or IFNβ-primed cells with DENV-2 for 24hrs and assayed viral genome replication by RT-qPCR (Table S5). Upon IFN pre-stimulation, wild-type cells showed dramatically reduced levels of DENV-2 RNA (122.5 fold reduction compared to unstimulated wild-type cells; Fig. S20), consistent with a strong IFNβ-mediated antiviral effect. In line with our experiments with SARS-CoV-2, IFNβ-treated IFNAR2-S KO cells showed the greatest protective effect of IFN (1.9x higher compared to untreated IFNAR2-S KO cells, Fig. 4E), with significantly less DENV-2 RNA measured compared to IFN-treated wild-type cells (p-value = 0.0029).

Combined, our viral infection experiments indicate that IFNAR2-S has a significant role in the antiviral response mediated by type I IFNs, and that the relative expression levels of IFNAR2-S and IFNAR2-L isoforms are likely to play an important role in tuning cellular responses to infection.

### IFNAR2-S evolved through a SINE-conversion event at the beginning of primate radiation

Finally, we investigated the evolution of the Alu-Jr exonization event that gave rise to the IFNAR2- S isoform. Based on a whole-genome alignment of 241 vertebrate species^54^ (Fig. S21), we determined that the intronic Alu-Jr element appeared ∼60-70 million years ago^55^ in simian primates (Fig. 5A). Interestingly, while the Alu-Jr element is not conserved in prosimian primates, we identified a more ancient Alu FAM element at the syntenic location in prosimian genomes that does not generate an exon (Fig. S22A). This observation suggests that the Alu-Jr element may have originated through a gene conversion event between similar Alu elements^56,57^ rather than a *de novo* retrotransposition event. Consistent with this scenario, the exonized Alu-Jr spans only the right monomer which shares sequence similarity to the original Alu FAM (Fig. S22B), while most Alu-Jr elements contain both a left and right monomer. The Alu-Jr element is oriented antisense to the *IFNAR2* gene in the last intron, which provides a splice acceptor site through its poly-A sequence^17^, which was a key step for the exonization of this element.

**Fig. 5.**
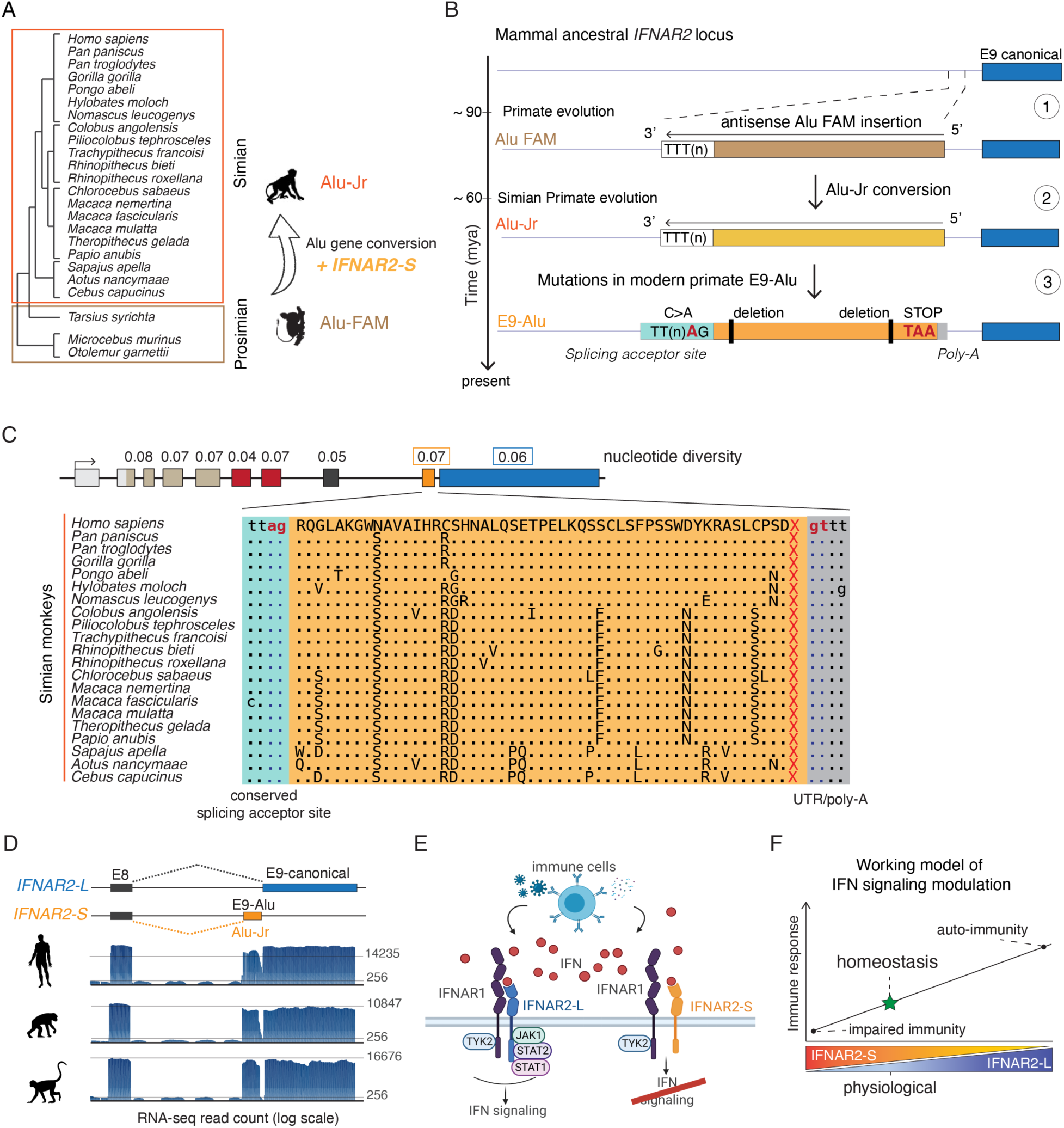
Evolution of the IFNAR2-S isoform and signaling. **A)** Primate phylogeny illustrates the evolution of IFNAR2-S at the time of evolution of simian primates ∼70-60 mya. **B)** Schematic reconstruction of IFNAR2-S evolution through the conversion of a syntenic ancestral FAM SINE by a simian primate-specific Alu-Jr Alu. Mutations in the Alu-Jr then allowed for its exonization and the evolution of IFNAR2-S. **C)** Sequence conservation of IFNAR2-S at the nucleotide (nucleotide diversity on top) and amino acid levels (alignment at the bottom). A dot represents a conserved amino acid. **D)** IFNAR2 exon coverage from NCBI aggregate RNA-seq data for three different simian primates. **E)** Model of IFNAR2 signaling in primates, where the type I IFN pathway is modulated by the presence of the decoy receptor IFNAR2-S, which binds IFN molecules but cannot initiate a signaling response. **F)**. Cellular immune homeostasis can be tuned by the IFNAR2-L : IFNAR2-S ratio.

We identified multiple additional mutations that likely facilitated the co-option of the Alu-Jr exon based on an alignment between the IFNAR2 Alu-Jr and the Alu-Jr consensus sequence. One C>A mutation 2bp upstream of the beginning of the E9-Alu exon introduced a stronger splicing acceptor site (AG instead of CG) (Fig. 5B and Figs. S22 and S23). Similarly, two T>A mutations may contribute to its escape from hnRNP and repressive splice factors^58^. To assess the weight of single nucleotide mutations on the Alu-Jr alternative splicing and exonization, we used the *Alubaster* software^59^ to perform an *in-silico* single nucleotide mutation screening. The *Alubaster* algorithm correctly identified the Alu-Jr of IFNAR2-S as exonized (base score of 0.59), and allowed for the identification of nucleotides that if mutated would negatively impact the exonization of the Alu-Jr, in particular at the splicing acceptor site (Table S6 and Fig. S23), and a SNP associated with severe Covid-19 (ID: 21:33262573 G>A) that is predicted to negatively affect the Alu exonization.

Within simian primates, the Alu-Jr exon is highly conserved, with an average nucleotide diversity of 0.07 (comparable to exons 4, 5 and 7, whereas the canonical exon 9 has a nucleotide diversity of 0.06). In addition, the Alu-Jr splicing acceptor site, coding sequence, stop codon, and 3’ untranslated region are conserved across all simian primates examined (Fig. 5C). Based on RNA-seq data, we confirmed that expression of the Alu-Jr exon and *IFNAR2-S* isoform is conserved across simian primates (Fig. 5D). The primate-wide conservation of both IFNAR2-S sequence and expression indicates that this Alu exonization event was co-opted to serve an essential role in regulating innate immune signaling.

## DISCUSSION

In this study, we show how TE exonization has shaped the evolution and regulation of the primate type I IFN signaling pathway. We demonstrate that alternative splicing of an Alu element within IFNAR2 generates the IFNAR2-S isoform, a novel decoy receptor that maintains immune homeostasis by regulating cellular sensitivity to type I IFNs (Fig. 5E and 5F). This work has several significant implications related to understanding IFN regulation, inflammatory disease mechanisms, and genome evolution.

First, our work reveals that alternative splicing of IFNAR2 and the expression of the IFNAR2-S isoform is a critical mechanism that regulates sensitivity to type I IFN signaling. Although the IFNAR2-S isoform was first identified more than 30 years ago, its potential physiological role has been largely overlooked and IFNAR2-L is assumed to be the only functionally relevant isoform. IFNAR2-S has been proposed to have dominant negative activity^29,60^, but the only functional study of IFNAR2-S to date was conducted using transfected mouse cells^31^, which lack IFNAR2-S and instead express rodent-specific IFNAR2 isoforms^61^. Our isoform-level dissection of IFNAR2-S and IFNAR2-L in multiple human cell lines reveals that IFNAR2-S is a potent decoy inhibitor of IFN signaling, broadly functioning to maintain immune homeostasis alongside other negative regulators of IFN including USP18, ISG15, and SOCS3^62–64^. While more work will be required to fully understand the mechanism of IFNAR2-S expression and function as an IFN signaling inhibitor, our work indicates that IFNAR2-S serves a critical role in dampening the activation of responses to type I IFNs.

Second, our work uncovers the relative expression of IFNAR2-L and IFNAR2-S as a potentially key determinant of the severity of immune-mediated diseases. Dysregulated IFN signaling underlies aberrant responses to infection, including severe COVID-19^45,47^, autoimmune diseases including type I interferonopathies such as systemic lupus erythematosus^65^, altered tumor immune phenotypes^66–68^ and is a hallmark of Down Syndrome^69,70^. Moreover, dysregulated IFN responses also contribute to acquired resistance to recombinant IFN therapy^71^. We speculate that interindividual variation in IFNAR2 isoform expression ratios –potentially due to non-coding disease variants– may drive differences in the severity of immune-related pathologies. Accordingly, the relative splicing of IFNAR2 isoform expression could represent a therapeutic target for modulating cellular sensitivity to infectious diseases or IFN-based therapies.

In agreement with early studies^72^, we find that both isoforms can form a ternary complex with IFNAR1 in the presence of IFN. However, assays for cellular IFNb uptake show that cells expressing only IFNAR2-L (IFNAR2-S KO) have consistently higher levels of internalized IFNb, whereas cells expressing only IFNAR2-S (IFNAR2-L KO) show overall lower levels of internalized IFNb. These differences may be explained by two alternative models: I) due to the fact that IFNAR2-S lacks the canonical intracellular recognition domains associated with endosomal transport^28^, the IFNAR2-S:IFN:IFNAR1 complex may not be internalized. Under this model, IFNAR2-S would act as a temporary trap decoy receptor for type I IFN molecules at the cell membrane. II) The IFNAR2-S:IFN:IFNAR1 complex may be less stable of, and therefore internalized at different rates than, the canonical ternary complex. Under this model, only a fraction of IFNAR2-S:IFNb:IFNAR1 complexes will be effectively internalized. III) Alternatively, the IFNAR2-S ternary complex may be internalized similarly to the IFNAR2-L canonical ternary complex, but IFN molecules may be recycled rather than degraded in the lysosomes through the canonical trafficking pathways^1^. Further studies will be required to clearly elucidate mechanisms of IFNAR2-S:IFNb:IFNAR1 trafficking.

Finally, our study demonstrates a novel example of a TE exonization event that was co-opted to serve a critical immune regulatory function. TE exonization is a widespread and well documented phenomenon in mammalian genomes, and several examples of adaptive TE exonization events have been documented^18,73,74^. However, it is broadly assumed that most events are nonfunctional because exonized TEs typically reside in untranslated regions, are subject to nonsense-mediated mRNA decay, or are actively repressed by the splicing machinery^21–23,58,75–77^. Our long-read transcriptomic analysis suggests that many immune genes express yet-uncharacterized isoforms generated by TE exonization, which may facilitate species-specific immune evolution. Our work implicates TE exonization as an underappreciated process that fuels the evolution of species-specific isoforms and the emergence of new mechanisms of immune regulation.

## Author Contributions

EBC and GIMP designed the study. GIMP, DMS, LLN, AI, HA, K. Gapin, IH performed experiments and analyzed data. K. Guo, QY, ABG, and CJW conducted viral infection assays. MLS, BGB, SLS, and EBC supervised the study. EBC and GIMP wrote the manuscript with input from all co-authors. All authors gave final approval for publication.

## Acknowledgements

We thank M. Good for technical assistance with cell culture. We thank the University of Colorado Genomics Shared Resource, Theresa Nahreini and the Flow Cytometry Shared Core Facility (NIH Grant *S10ODO21601*), and BioFrontiers Computing core for technical support during this study.

## RESOURCE AVAILABILITY

### Lead contact

Further information and requests for resources and reagents should be directed to and will be fulfilled by the lead contact, Edward B. Chuong.

### Materials availability

Any additional information required to generate newly generated materials reported in this paper is available from the lead contact upon request.

### Data and Code availability

A UCSC genome browser session with data generated for this study is available at: https://genome.ucsc.edu/s/GiuliaPasquesi/HeLa_KO_4hIFNb.

RNA-seq sequencing reads generated for HeLa wild-type and KO cells are deposited in NCBI GEO Accession: GSE211812

### Supplementary Materials

Figs. S1 to S23

Tables S1 to S7

References (*1*–*96*)

## METHODS

### Re-analysis of publicly available RNA-seq datasets

#### GTEx long-read

The FLAIR transcriptome *gtf* file (long-read based annotation) generated by the GTEx consortium was downloaded from https://www.gtexportal.org/home/datasets. Cufflinks v2.2.1 ^78^ (*gffread* function) was used to convert the FLAIR *gtf* to *fasta* format, and Salmon v1.10.0^79^ was used to create a salmon index from the resulting FLAIR *fasta* file. GTEx V9 Protected Access long-read RNA-seq *fastq* files (92 samples total; https://gtexportal.org/home/protectedDataAccess) were downloaded using AnViL (https://anvil.terra.bio/). *Fastq* file names were modified to add the tissue type for each sample. Minimap2 v2.22-r1101 ^80^ was used to align each *fastq* file to the FLAIR transcriptome *fasta* file (parameters *-a -t 4*). Samtools v1.16.1 ^81^ was used to convert *sam* files into *bam* format and sort by read name. Salmon was used with parameters: *--ont --libType A -p 8 --rangeFactorizationBins 4 -w 10000* and the FLAIR *fasta* and *gtf* files to quantify isoforms.

We then extracted exon coordinates from the FLAIR *gtf* file and intersected them with the Dfam transposable element human annotation (hg38 RepeatMasker open-4.0.6 - Dfam 2.0 ^82^), filtering for features where more than 80% of a TE overlapped an annotated exon. To focus on TEs that provide exon splice donor or acceptor sites, and to exclude TEs that localize to untranslated regions (UTRs), we generated a custom splice-site *gtf* file based on the exon coordinated of the FLAIR *gtf* file, and used this to further refine TE-exonization events. We then retained only entries corresponding to protein coding genes and that had a transcriptional support of TPM > 5 in at least one GTEx V9 Protected Access long-read RNA-seq sample. We selected candidate isoforms of genes involved in immune and inflammatory responses by filtering the total list of TE-derived isoforms for GO terms: "Inflammatory Response" (GO:0006955), "Immune Response" (GO:0006954), "Response to Cytokine" (GO:0034097), Cytokine Activity (GO:0005125), "Cytokine production" (GO:0001816), "Regulation of Cytokine Production" (GO:0001817), "type I interferon production" (GO:0032606), "type I interferon signaling pathway" (GO:0060337), and "response to type I interferon" (GO:0034340). All results are reported in Supplementary Table 1.

To analyze isoform-specific expression of IFNAR2, we generated a custom FLAIR *gtf* file to only include two possible IFNAR2 isoforms: ENST00000342136.8 (*IFNAR2-L*) and ENST00000404220.7 (*IFNAR2-S*). Other, spurious IFNAR2 annotations were removed. The above minimap2 and salmon quantification steps were then repeated using this modified FLAIR *gtf* and *fasta* file. The two isoforms of interest were extracted from each sample, collated and plotted.

#### Macrophage long-read cDNA sequencing

To analyze the contribution of transposable elements (TEs) to isoform diversification in the context of immune responses, we analyzed paired short-read Illumina RNA-seq and long-read Nanopore cDNA sequencing data from a monocyte-derived macrophage stimulation panel ^26^. We used the long-read sequencing assembled isoforms alignment *bam* file available from the UCSC genome browser (https://genome.ucsc.edu/s/vollmers/IAMA) to generate a *gff* file that included exon boundary coordinates ^83^. We then intersected exons coordinates with the Dfam transposable element human annotation (hg38 RepeatMasker open-4.0.6 - Dfam 2.0 ^82^), filtering for features where more than 80% of an exon overlapped a TE. Candidate isoforms were finally subset based on their biological function, as done for the GTEx V9 Protected Access long-read dataset and visually inspected to annotate TE exonization events as in protein coding or UTR regions (Table S1).

#### GTEx and TCGA short-read

We assessed expression levels of *IFNAR2-L* and *IFNAR2-S* by querying data for multiple tissues and individuals using the GTEx portal repository (http://gtexportal.org) ^32^ and the Gepia2 portal (http://gepia2.cancer-pku.cn/) ^33^, which leverages TCGA and GTEx resources. We downloaded pre-quantified and normalized transcript and exon read counts, as well as normalized exon-exon junction read counts from the GTEx analysis V8 (dbGaP Accession phs000424.v8.p2). We used the following ENSEMBL transcript IDs for each isoform: *IFNAR2-L*: ENST00000342136 and ENST00000683941; *IFNAR2-S*: ENST00000404220 and ENST00000382264. For matched healthy and tumor sample isoform expression quantification, we used the Gepia2 portal web interface to access isoform count data. We matched these short-read RNA-seq datasets with long-read data from the GTEx portal (https://gtexportal.org/home/datasets; GTEx Analysis V9). For the long-read analysis, we used the following HAVANA transcript IDs for each isoform: *IFNAR2-L*: OTTHUMT00000139825.1; *IFNAR2-S*: OTTHUMT00000139824.4 and OTTHUMT00000139827.1

#### Viral infection datasets

Changes in relative isoform expression in human cell lines upon viral infections were examined using paired-end read data only, when high read coverage and quality, as well as biological replication level was available. We used Salmon v1.8.0 ^79^ to map reads and quantify transcript expression levels (TPM) from studies of SARS-CoV2 infected Calu3 cells ^43^ and Influenza B infected A549 cells ^44^. Salmon was run using the following parameters: *salmon quant --libType A --validateMappings --rangeFactorizationBins 4 --gcBias -w 10000*. We used the hg38 assembly of the human genome and the latest available Gencode ^84^ comprehensive release for the human transcriptome (release 40; GRCh38.p13) as references.

#### Non-human primates

Aggregate RNA-seq exon read coverage from multiple tissues were obtained from NCBI annotations of *Homo sapiens*: https://www.ncbi.nlm.nih.gov/gene/3455; *Pan troglodytes*: https://www.ncbi.nlm.nih.gov/gene/473965; and *Macaca mulatta*: https://www.ncbi.nlm.nih.gov/gene/699726.

### Evolutionary analysis of Alu-Jr

The conservation of the Alu-Jr element was analyzed with respect to the 241-way Vertebrate Genome Alignment in the UCSC genome browser. Alignment of CDS and protein sequences of IFNAR2-S and IFNAR2-L for 21 primate species was manually curated in AliView ^85^, and used as reference to calculate nucleotide divergence using MEGA v.X ^86^. Mutations that occurred in the human E9-Alu exon compared to Alu-Jr and Alu-FAM consensus sequences were manually detected based on the alignment generated using the Dfam website (dfam.org ^87^).

### Disease variant analysis of IFNAR2

The COVID-19 Host Genetics Initiative data browser was used to visualize variants associated with hospitalized COVID-19 patients at the IFNAR2 locus (chr21:32982032-33503778; Release: R6; Phenotype: B2_ALL: Hospitalized COVID19+ vs. population control: Analysis: r6-nStudies- 43-nCases-24274-nControls-2061529 (leave_23andme)). Severe COVID variants were visualized using the "COVID GWAS v4" track on the UCSC hg38 genome assembly (Severe respiratory COVID risk variants from the COVID-19 HGI GWAS Analysis A2 (4336 cases, 12 studies, Rel 4: Oct 2020), with a minimum -log10 p-value = 3, Effect size range -1.6 to 2.2.

### Cell culture

Experiments were conducted in validated HeLa (ATCC CCL-2; RRID:CVCL_0030), A549 (ATCC CCL-185; RRID:CVCL_0023), and PEO1 (RRID:CVCL_2686; Gynecologic Tumor and Fluid Bank, University of Colorado Anschutz) cell lines. All cell lines were grown at 37°C and 5% CO_2_. HeLa cells were grown in DMEM (Thermo Fisher #10-565-018) supplemented with 10% fetal bovine serum (FBS) and 100U/mL Penicillin-Streptomycin (P/S). A549 cells were grown in F12- K nutrient mixture (Thermo Fisher #21-127-022) supplemented with 10% FBS and 100U/ml P/S. PEO1 cells were grown in RPMI 1640 media (Gibco #11875085) supplemented with 10% FBS and 100U/ml P/S. Cells were periodically tested for the presence of mycoplasma using the ATCC Universal mycoplasma detection kit (#30-1012K). Briefly, 10^4^-10^5^ adherent cells were scraped in their culturing conditions and lysed following manufacturer’s instructions. Provided universal primers were used to amplify genomic DNA from sample cells and from a positive control (pUC19:: *M.arginini*).

### Cell treatments

Cells were seeded in either 10cm dishes, 6 well plates or 24 well plates the day before treatment, and left in incubation overnight. For IFNβ experiments, 10U/ml of animal-free recombinant human IFNβ (Proteintech #HZ-1298) were used as standard dose, unless otherwise specified. For RT-qPCR and RNA sequencing, cells were treated for 4hrs. For protein extraction and assessment of phosphorylated STAT1/2 and STAT1/2, cells were treated for 30min. For viability assays (CellTiter Glo and Crystal Violet), cells were treated for 4 days without media replacement. For control/mock-treated cells, equal volumes of DPBS were used instead of IFNβ. For the cytokine panel assay, the following cytokines were used at a 100U/ml: i) Human IFNα2 (Proteintech #HZ- 1066) and ii) Human IFNγ (R&D systems #285-IF-100/CF). For HaloTag protein pull-down experiments, cells were grown in 15cm dishes and treated with 1000U/ml of IFNβ for 5min or 1000U/ml of IFNα2 for 10min.

### HiBiT epitope tagging

To detect the presence of IFNAR2-S proteins in HeLa cells, we followed a CRISPR knock-in protocol to endogenously insert a HiBiT tag (Promega ^88^) at the 3’ terminus of the *IFNAR2-S* mRNA (Fig. S6A). Briefly, wild-type HeLa cells were electroporated with the IDT Alt-R CRISPR-Cas9 system (recombinant Cas9, tracrRNA, crRNA) with a single-stranded oligodeoxynucleotides (ssODN) donor template and a guide RNA (gRNA) targeting the 3’ end of the IFNAR2-S E9-Alu (Table S7). The ssODN template was designed to append a terminal HiBiT tag upstream of the endogenous stop codon. Both the ssODN and gRNA were synthesized by IDT. Clonal lines were isolated using the limited dilution method in a 96 well plate format, and heterozygous and homozygous clones were identified and screened using the Nano-Glo HiBiT Lytic Detection System (Promega #N3030). We confirmed in-frame insertions by PCR, Sanger sequencing and western blot (Fig. S6).

### CRISPR deletions

We generated CRISPR-mediated knockout of *IFNAR2-S*, *IFNAR2-L*, and *IFNAR2*-exon 7 (IFNAR2 KO) knockout HeLa cell lines (Fig. 2A). For each knockout, two gRNAs were identified using the CRISPR targets track for GRCh38/hg38 release of the human genome on the UCSC genome browser (Table S7). Guides were synthesized by Integrated DNA Technologies (IDT). HeLa cells were electroporated with the IDT Alt-R CRISPR-Cas9 system (recombinant Cas9, tracrRNA, crRNA) and paired gRNAs using the Neon system (ThermoFisher Scientific). Clonal lines were isolated using the limited dilution method in a 96-well plate format, and single clones were identified and screened for homozygous deletions by PCR using both flanking and internal primer pairs at the expected deletion site (Fig. S4A). The same approach was used to generate A549 *IFNAR2-S* and *IFNAR2-L* knockout cell lines (Fig. S18).

### Small Interfering RNAs

To conduct IFNAR2 isoform-specific depletion by small interfering RNA (siRNA), we designed two siRNA oligos per target exon (Table S7). Custom siRNAs were purchased from Horizon Discovery and modified with the ONTARGET-plus modification. The day before transfection, cells were seeded in 24 well plates and allowed to grow overnight. According to the manufacturer’s protocol, siRNAs were diluted to a concentration of 5μM in 1x siRNA buffer (Horizon Discovery #B-002000- UB-100), and transfected at a final concentration of 25nM per well (5μM siRNA) using the DharmaFECT 1 Transfection reagent (Horizon Discovery #T-2001-02). Cells were left in incubation for 6hrs, after which transfection media was replaced with regular growing media. We used a positive control (siGLO Lamin A/C Control siRNA; Horizon Discovery #D-001620-02-05) to visually assess transfection efficacy. Transfection efficiency was also assessed through RT-qPCR for the *LMNA* gene. We used ON-TARGETplus Non-targeting Control siRNA #2 as a negative control siRNA (Horizon Discovery #D-001810-02-05). To assess whether isoform-specific modulation of IFNAR2 expression affects cellular responses to IFN treatment, we additionally performed a titration experiment by co-transfecting the non-targeting control siRNA and target siRNAs for a total molarity of 10μM in the following combinations: 0μM target + 10μM negative control; 2μM target + 8μM negative control; 4μM target + 6μM negative control; 6μM target + 4μM negative control; 8μM target + 2μM negative control; 10μM target + 0μM negative control.

### Generation of stable overexpression cell lines

#### IFNAR2-S overexpression

The *IFNAR2-S* coding sequence (CDS) was synthesized by Twist Bioscience. We included a 5’ handle, *NheI* restriction site and Kozak consensus sequence at the 5’ terminus upstream of the first codon (ATG) and a *NotI* restriction site and 3’ handle downstream of the stop codon (TGA). Fragments were resuspended at a final concentration of 100ng/ul in EB buffer and then cloned into a PB-CMV-MCS-EF1α-Puro PiggyBac cDNA Cloning and Expression Vector (PB vector hereafter, System Biosciences #PB510B-1). Following ligation, PB vectors were transfected in NEB stable competent *E. coli* cells (New England Biolabs #C3040H). Transfected cells were grown for 24hrs at 30°C on agar plates treated with LB/Carb100. From each plate, 3 colonies were randomly selected and further expanded for plasmid DNA purification (Zyppy™ Plasmid Purification Kit; Zymo research #D4019). Extracted DNA was then sequenced to verify the integrity of the cloned fragments.

#### ACE2 overexpression

The ACE2 coding sequence was synthesized by PCR amplification from the pUC57-ACE2 plasmid (GenScript). We included a *NheI* restriction site at the 5’ terminus and a *PmeI* restriction site at the 3’ terminus. The ACE2 fragment was digested with *DpnI* overnight to remove plasmid template from the PCR sample before cloning into the expression vector pB-CAGGS-dCas9-KRAB-MeCP2. Final expression vector was prepared by restriction digest to remove the dCas9-KRAB-MeCP2 domains using the restriction enzymes *PmeI* and *NheI*. The pB-CAGGS vector and ACE2 insert were ligated by In-Fusion cloning (Takara Bio #638947) following manufacturer’s instructions. Following ligation, pB-CAGGS-ACE2 vectors were transfected in NEB stable competent *E. coli* cells (New England Biolabs #C3040H). Transformed cells were grown for 24hrs at 30°C on agar plates treated with 100μg/ml carbenicillin. From each plate, 3 colonies were randomly selected and further expanded for plasmid DNA purification (E.Z.N.A. Plasmid DNA Mini Kit I, Omega #D6942-02). Extracted DNA was then sequenced to verify the integrity of the cloned fragments. Colonies that had integrated the original fragments were then further expanded, and plasmid DNA was extracted by using the Zymopure II plasmid midiprep kit (Zymo research #D4201).

#### IFNAR2-S-HaloTag and IFNAR2-L-HaloTag overexpression

IFNAR2-L and IFNAR2-S coding sequences were cloned from previously generated piggybac vectors using a PCR amplification protocol, and then cloned into the pHTC HaloTag CMV-neo Vector (Promega #G7711) using In-Fusion cloning (Takara Bio #638947). For the generation of empty HaloTag expressing vectors, the HaloTag coding sequence was amplified from the pHTC HaloTag CMV-neo Vector (Promega #G7711). Cloning reactions were transfected into NEB stable competent *E. coli* cells (New England Biolabs #C3040H) and grown overnight at 37°C on agar plates treated with 100μg/ml carbenicillin. Plasmids were extracted from 3 bacterial colonies each using the E.Z.N.A. Plasmid DNA Mini Kit I (Omega #D6942-02) and sequence verified using Plasmidsaurus long-read sequencing (https://www.plasmidsaurus.com/).

For each plasmid, colonies that were sequence-validated to have integrated the original fragments were then further expanded, and plasmid DNA was extracted by using the Zymopure II plasmid midiprep kit (Zymo research #D4201). Transfection of plasmid DNA (2.5μg vector + 1μg of transposase plasmid) was performed using the FuGENE transfection reagent (Promega #E5911) according to the manufacturer’s instructions. Cells were selected for the integration of the plasmid DNA using Puromycin (2μg/ml) or Blasticidin (5μg/ml) for 3 to 4 days and validated by RT-qPCR.

### Assessment of ISG and IFNAR2 isoform expression levels by RT-qPCR

Cells were lysed in 300μl of RNA lysis buffer (Zymo Research #R1060-1-50), and stored at -80°C until RNA extraction was performed using the Quick-RNA MiniPrep kit (Zymo Research #R1054) following manufacturer’s instructions. A NanoDrop One spectrophotometer (Thermo Fisher Scientific) was used to determine RNA concentration and quality; all samples passed quality assessment. For cytokine treatments and Dengue viral genome replication experiments, 3 experimental replicates per condition were analyzed; siRNA experiments were independently repeated twice. For analysis of ISG expression and validation of IFNAR2-L/S knockout, three different HeLa CRISPR/Cas9 clones were used. One clone per genotype was then chosen for further experiments (i.e., cytokine panel, transfection of overexpression constructs, Dengue viral infection). A list of primers used for RT-qPCR and cycling conditions are provided in Supplementary Table 7. RNA expression levels were quantified using the Luna Universal One-Step RT-qPCR Kit (New England Biolabs #E3005L) according to the manufacturer’s instructions. In brief, for each reaction 25ng of RNA was combined with 5μl 2× Luna Universal One-Step Reaction Mix, 0.5μl 20× Luna WarmStart RT Enzyme Mix, 0.4μl 10μM forward primer, and 0.4μl 10μM reverse primer. Reactions were amplified using a CFX384 Touch Real-Time PCR Detection System (Bio-Rad). On-target amplification was assessed by melt curve analysis. Each sample was run either in technical duplicate or triplicate. RT-qPCR result values were analyzed using the ΔCq expression method (2^(-ΔCq)) normalizing Ct values of target genes to the Ct value of the *CTCF* housekeeping gene per each sample/replicate. Statistical significance for the treated vs. control comparisons was assessed using either a two-tailed t-test in R (*t.test* function) or a two-tailed Mann-Whitney U Test (*wilcox.test* function) upon assessment of normality of the data (*shapiro.test* function) and that other assumptions were not violated. To compare the effect of different levels in multifactorial experimental designs (i.e., the effect of treatment across cell lines), a multifactor analysis of variance (ANOVA) was performed in R upon ΔCq expression normalization of treated samples by the average of the untreated (*contrast(emmeans(model, ∼factor), interaction = "pairwise")* function).

### RNA-seq library preparation of wild-type and KO HeLa cells

RNA from control and 4hrs IFNβ treated HeLa wild-type and CRISPR/KO cells was extracted as described above. PolyA enrichment and library preparation was performed using the KAPA mRNA HyperPrep Kit (Kapa Biosystems #8098115702) according to the manufacturer’s protocols. Briefly, 500ng of RNA was used as input, and single-index adapters (Kapa Biosystems #08005699001) were added at a final concentration of 10nM. Purified, adapter-ligated library was amplified for a total of 11 cycles following the manufacturer’s protocol. The final libraries were pooled and sequenced on an Illumina NovaSeq 6000 (University of Colorado Genomics Core) to obtain ∼20million 150bp paired-end reads per library.

### RNA-seq analysis of wild-type and KO HeLa cells

Paired-end 150bp read length fastq files were quality-filtered and adapter trimmed using BBDuk v.38.05 ^89^; quality check was performed using FastQC v0.11.8 ^90^ and inspected through MultiQC v1.7 ^91^. Filtered fastq files were then mapped to the human hg38 genome using a 2-pass approach in STAR v2.7.3a ^92^. STAR was run following default parameters and allowing for multi-mapping reads (with options ‘*–outAnchorMultimapNmax 100 –winAnchorMultimapNmax 100 – outFilterMultimapNmax 100*’). The latest Gencode main comprehensive annotation of the human genome available at the time of analysis (release 40; GRCh38.p13) was used as reference for the mapping process. For the second pass of mapping, we filtered out novel junctions that mapped to the mitochondrial genome. Resulting alignment files in sorted *bam* format were then provided as input for gene expression quantification using featureCounts ^93^. Pairwise differential expression analysis for each genotype was performed in DESeq2 v1.32 ^94^. To assess the difference in treatment effect on gene expression between wild-type and IFNAR-S KO cells we used wild-type and untreated conditions as reference levels in the experimental design: *dds <- DESeqDataSetFromMatrix(countData = val, colData = coldata, design = ∼ genotype + treat + genotype:treat)*. Specifically, we tested for the treatment effect on IFNAR2-S KO cells using the formula: *results(dds_ds, list(c("treat_IFN_vs_cnt","genotypeKO.treatIFN")))*, as advised in the DESeq2 manual. Functional enrichment analyses of differentially expressed genes (adj. *p*-val < 0.05, log2FC > 2 were performed using the WebGestalt web tool ^95^.

### Protein detection by SDS-PAGE

Proteins were extracted from pelleted cells using RIPA buffer (+ 1x RPMI and 1x protease inhibitor). Briefly, cell pellets were washed twice with ice-cold DPBS, and then resuspended in RIPA buffer (∼50-100μl per million cells). Lysates were then incubated on ice for 15min, followed by sonication and a final incubation on ice for additional 15min. Cellular debris were pelleted by centrifugation (13,000g at 4°C for 5min). The supernatant, containing both membrane and cytosolic proteins, was collected and protein quantified using the BCA assay (Thermo Scientific #23227). Samples were denatured in 1x Licor loading buffer (LI-COR Biosciences #LIC-928- 40004) and 1:10 β-mercaptoethanol (BME) at 70°C for 10min. Antibody-based Western blotting was performed following a SDS-PAGE protocol optimized for detection using the LI-COR Odyssey CLx Infrared Imaging System. A list of antibodies used is provided in Table S7. To detect the HiBiT-tagged IFNAR2-S isoform, we used the Nano-Glo HiBiT Blotting System (Promega) to detect protein following SDS-PAGE and transfer. We added Nano-Glo Luciferase Assay Substrate and LgBiT protein (Promega) and imaged using the ImageQuant LAS4000 Imager.

### Phospho-Flow Cytometry

To quantify levels of phosphorylated STAT1 in wild-type and knockout HeLa cells, cells were seeded in 10cm dishes and grown until 70% confluent. The following day, cells were treated with different concentrations of IFNβ (0U/ml, 10U/ml, 100U/ml, 1000U/ml and 2000U/ml) for 30min. Media was then removed and cells washed twice in ice-cold DBPS. A cell scraper was used to lift the cells. Cells were fixed (BD Biosciences 554655) for 10 min at 37°C, and permeabilized (BD Biosciences 558050) for 30min at 4°C. Cells were then stained with a PE-labeled anti pY701- STAT1 Ab (BD Biosciences 612564) and analyzed using an Accuri C6 Plus flow cytometer (BD Biosciences). Using FloJo V10.9 (BD Life Sciences), events were gated by singlets and cells identification, and median fluorescence intensity (MFI) was calculated for the whole gated population.

### HaloTag pull-down and protein detection

HeLa cells stably expressing IFNAR2-S or IFNAR2-L C-terminal HaloTag were grown in 15cm dishes and treated when 80% confluent (∼20M cells per plate). Upon treatment, cells were washed twice with ice-cold DPBS, and brought in suspension by scraping in 15ml of ice-cold DPBS. We set aside 1ml of cell suspension for total protein lysate validation, and used the remaining 14ml of cell suspension for the HaloTag pull-down. Cells were pelleted by centrifugation at 4°C for 5min at 2,000g, and cell pellets were stored at -80°C. We used the HaloTag Mammalian Pull-Down and Labeling Systems kit (Promega; #G6500) to identify protein interacting with IFNAR2 in stimulated and unstimulated conditions following manufacturer’s instructions. Briefly, cells were lysed using 300μl of mammalian lysis buffer and 6μl of 50x protease inhibitor cocktail, and left in incubation on ice for 5min. To reduce lysate viscosity, cells were passed through a 25- gauge needle 5 times, and lysates were cleared by centrifugation at 14,000g for 5min at 4°C. Diluted clear lysates were then bound to equilibrated HaloTag resin at 4°C for 2hrs on a tube rotator. We separated the bound and unbound fraction by centrifugation, and the bound fraction was washed 4 times following the manufacturer’s instructions. To release the bait protein tagged with the HaloTag and its protein partners, we used the ProTEV Plus enzyme (Promega; #V6101) to cleave the HaloTag from the bait protein following the protocol provided with the HaloTag Mammalian Pull-Down and Labeling Systems. For each step (pre-binding total lysate, unbound, wash and pull-down), a fraction was saved for optimization and validation purposes. Protein extracts were quantified as previously described.

### Immunofluorescence

Approximately 20,000 wild-type, IFNAR2-S and IFNAR2-L KO A549 cells were seeded in SensoPlates Plus 96 well plates (Greiner Bio-One; #655891) and grown in complete F12-K media until they reached 80% confluency. Cells were serum-starved for 4hrs before treatment with 0U/ml, 10U/ml and 1000U/ml of IFNβ for 5min, then washed once with DPBS. Cells were fixed in 4% paraformaldehyde for 15min, and washed three times for 5min with DPBS before staining. Cells were incubated with Wheat Germ Agglutinin (5μg/ml; Invitrogen #W11261) for 10min to stain cell membranes, then washed three times with DPBS before permeabilization in ice cold Phosflow Perm Buffer III for 10mins at -20°C (BD Bioscience #558050). Before incubation with primary antibodies, cells were blocked for 1hr at room temperature in 5% goat serum-based buffer (1x DPBS, 5% goat serum, 0.3% Triton X-100). Incubation was performed overnight at 4°C with 1:100 primary antibody dilution in 1% BSA-based buffer (1X DPBS, 1% BSA, 0.3% Triton X-100). The following day cells were washed three times with DPBS, and incubated for 1hr in dark condition at room temperature with the secondary antibody (table S7) at a dilution of 1:1000 in 1% BSA-based buffer. Following wash of the secondary antibody, cell nuclei were stained with 1ug/ml of DAPI for 10min. Cells were imaged using a Nikon Spinning Disc Confocal Yokogawa CSU X1 microscope using three different lasers (intensity: 488 nm (20%), 405 nm (15%), 640 nm (20%); EM Gain 10MHz, exposure: 488 nm (300ms), 405 nm (300ms) and 640 nm (600ms)) controlled by the NIS Element v5.42.03 software. For each channel, each image file comprised a z-stack of images, where images were taken at 0.5μm intervals over a total of 3μm in the z-axis. Each z-stack was condensed to one image based on the ‘maximum projection feature in the NIS Element software. Images were analyzed using the software Imaris x64 (v 10.0.0 ^96^), using the cell function. In brief, the nuclei were detected and segmented using the DAPI channel. The cell boundaries were determined using the signal intensity threshold method. The cell boundaries for STAT2 analysis were determined using the STAT2 signal in the Cy5 channel, and for the pSTAT2 analysis the Wheat Germ Agglutinin (wga) stain in the GFP channel was used. All cells that had inaccurate cell boundaries were removed manually from the analysis. To measure the nucleus:cytoplasmic ratio of STAT2 signal, the intensity of the signal in the Cy5 channel for each nucleus and corresponding cytoplasmic area in the nucleus was calculated per cell.

A similar approach was followed to image A549 cell lines expressing IFNAR2-HaloTag isoforms. For HeLa cell lines stably expressing IFNAR2-HaloTag isoforms no serum-starvation was required. Cells were treated with 100U/ml of IFNβ for 30min, and stained for 1hr at room temperature with the permeable Janelia Fluor HaloTag Ligand (JFX549; Promega #GA1110).

### Cell viability assays

All assays of ISG expression levels were paired with analyses of cell viability according to the conditions described above (*Cell Treatment* paragraph). We performed visual and quantitative assessment of cell viability upon treatment. We used crystal violet to fix and stain cells grown below confluency; crystal violet was then solubilized using a solution of DPBS, 10% acetic acid and 10% methanol. For each sample, absorbance at a wavelength of 562nm was read in triplicates. We also assessed cell viability of each sample in triplicates through a luminescence assay of ATP (CellTiter-Glo 2.0 Assay, Promega #G9242) following manufacturer’s instructions.

### Viral infections

#### SARS-CoV-19 infection

Wild-type and KO A549 stably expressing ACE2 were infected with the B.1.617.2 (BEI cat# NR-55672) strain of the SARS-CoV-2 virus following the methodology described in ^39^. Briefly, to assess viral replication levels an RT-qPCR approach was followed. 2.5×10^4^ cells per well were plated in a 48-well and pre-stimulated for 18hrs with IFNβ at a concentration of: 20pM, 2pM, 0.2pM, and 0pM (experiment 1) and 200pM, 20pM, 2pM, 0.2pM, 0.02pM and 0pM (experiment 2 and 3 to achieve more accurate IC_50_ estimation). Cells were then infected with 1.74×10^5^ PFU/well (1.57×10^8^/ul N1 copies; experiment 1) and 2.9×10^5^ PFU/well (2.61×10^8^/ul N1 copies; experiments 2 and 3) for 2hrs. Viral concentration was optimized upon experiment 1 to achieve a 10^5^ N1 yield in RT-qPCR detection for control samples that were not pre-stimulated with IFNβ. After two washes with PBS, 500μl complete media containing the corresponding concentrations were added. 24hrs post infection supernatants were harvested for RNA extraction and qPCR analysis. Total RNA was extracted from 100μl of cell culture supernatant using the E.Z.N.A Total RNA Kit I (Omega Bio-Tek; #M6399-01) and eluted in 50μl of RNAse-free water. 5μl of this extract was used for qPCR. Official CDC SARS-CoV-2 N1 gene primers (Supplementary Table 7) and TaqMan probe set were used with the Luna Universal Probe One-Step RT-qPCR Kit. The real-time qPCR reaction was run on a Bio-Rad CFX96 real-time thermocycler under the following conditions: 55°C 10mins for reverse transcription, then 95°C for 1min followed by 40 cycles of 95°C 10s and 60°C 30s. The absolute quantification of the N1 copy number was interpolated using a standard curve with 10^7^-10^1^ serial 10-fold dilution of a control plasmid (nCoV-CDC-Control Plasmid, Eurofins). For each experiment, each sample was read in triplicate or 4 times. The Bio-Rad CFX manager software was used to automatically calculate the absolute N1 copy numbers based on the standard curve generated from the seven 10-fold serially diluted standards (in triplicates) on each qPCR plate. Statistical analyses were performed in R as previously described.

#### Dengue infection

Wild-type and KO HeLa cells were seeded in 24 well plates at a seeding density of 80,000 cells/well in complete DMEM 24hrs before pre-treatment with IFNβ and incubated in regular conditions. Media was removed and replaced with complete media containing 100U/ml of IFNβ and incubated for 24hrs before infection. For infection, DENV-2 strain 16681 was used to prepare inoculum equivalent to a multiplicity of infection (MOI) = 1. Media was removed and monolayer was washed once with PBS, and 100 or 200μl of viral suspension diluted in PBS containing calcium and magnesium was added per 24 or 12 well plates, respectively. Cells were incubated with virus inoculum for 1hr at 37°C, with manual rocking every 15min; inoculum was removed, washed with PBS and complete media added for incubation in regular conditions. Supernatant was collected for further virus titration and cells lysed for RNA extraction (see *Assessment of ISG and IFNAR2 isoform expression levels by RT-qPCR* paragraph). Four experimental conditions were included: i) untreated control cells; ii) cells infected with DENV-2; iii) cells pre-treated with IFN; iv) cells pre-treated with IFN and infected with DENV-2.

