## Supplemental Figures for "Regulation of human interferon signaling by transposon exonization"

Supplementary Figures

Supplementary Figure 1

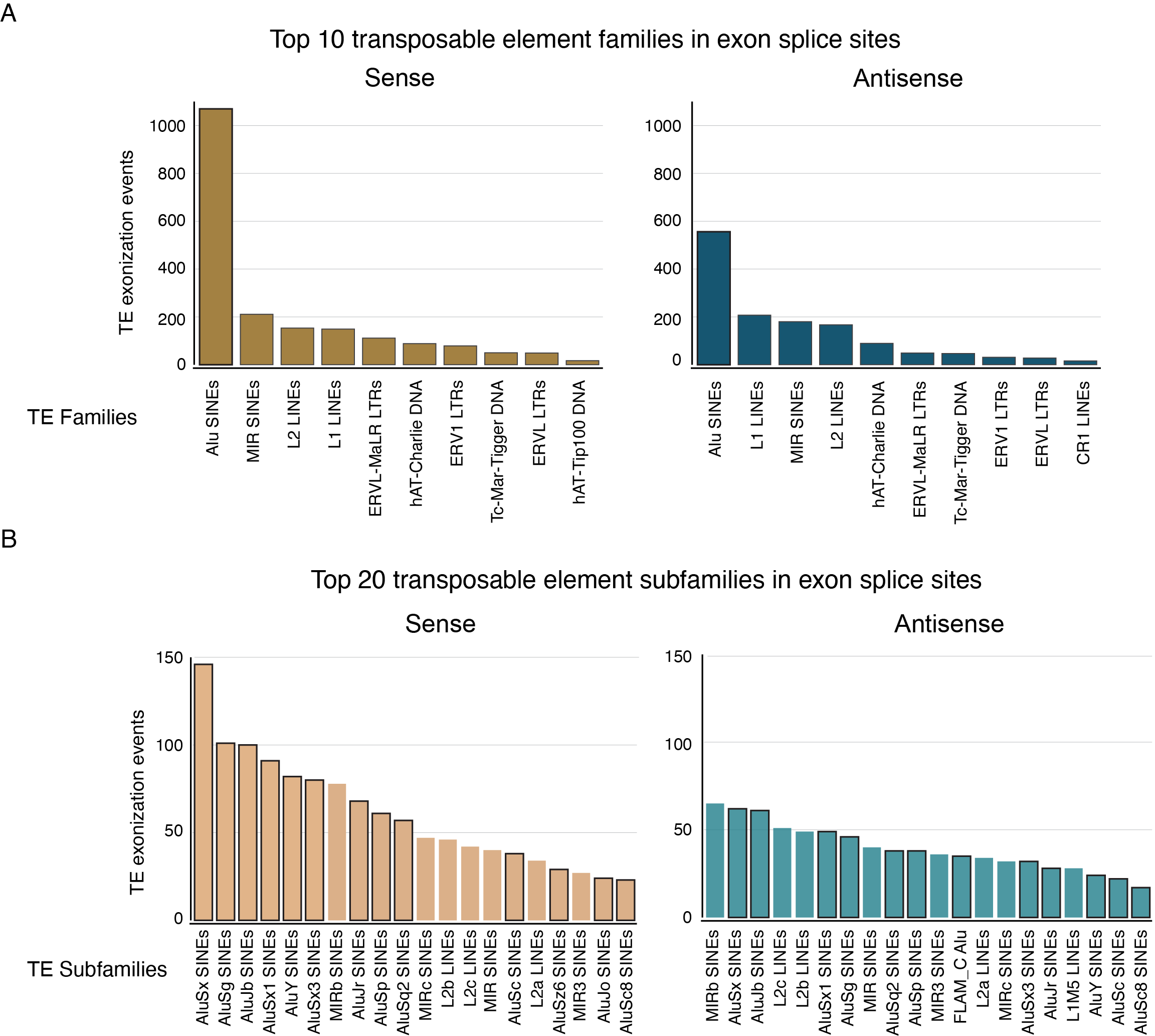

**Fig. S1) Most common transposable elements providing alternative exon splice sites.** TE exonization event counts for the top 10 TE family **(A)** and the top 20 TE subfamilies **(B)** based on their orientation compared to that of the protein-coding gene they provide the alternative splice site to (sense= same orientation, to the left; antisense = opposite orientation, to the right). Only TEs that overlapped a protein-coding gene exon for 80% of their length and a splice site for isoforms with TPM >5 in at least one sample were included (*27*). In (**B**) black outlines highlight Alu SINE transposable elements, the most abundant family of TEs to undergo exonization.

Supplementary Figure 2

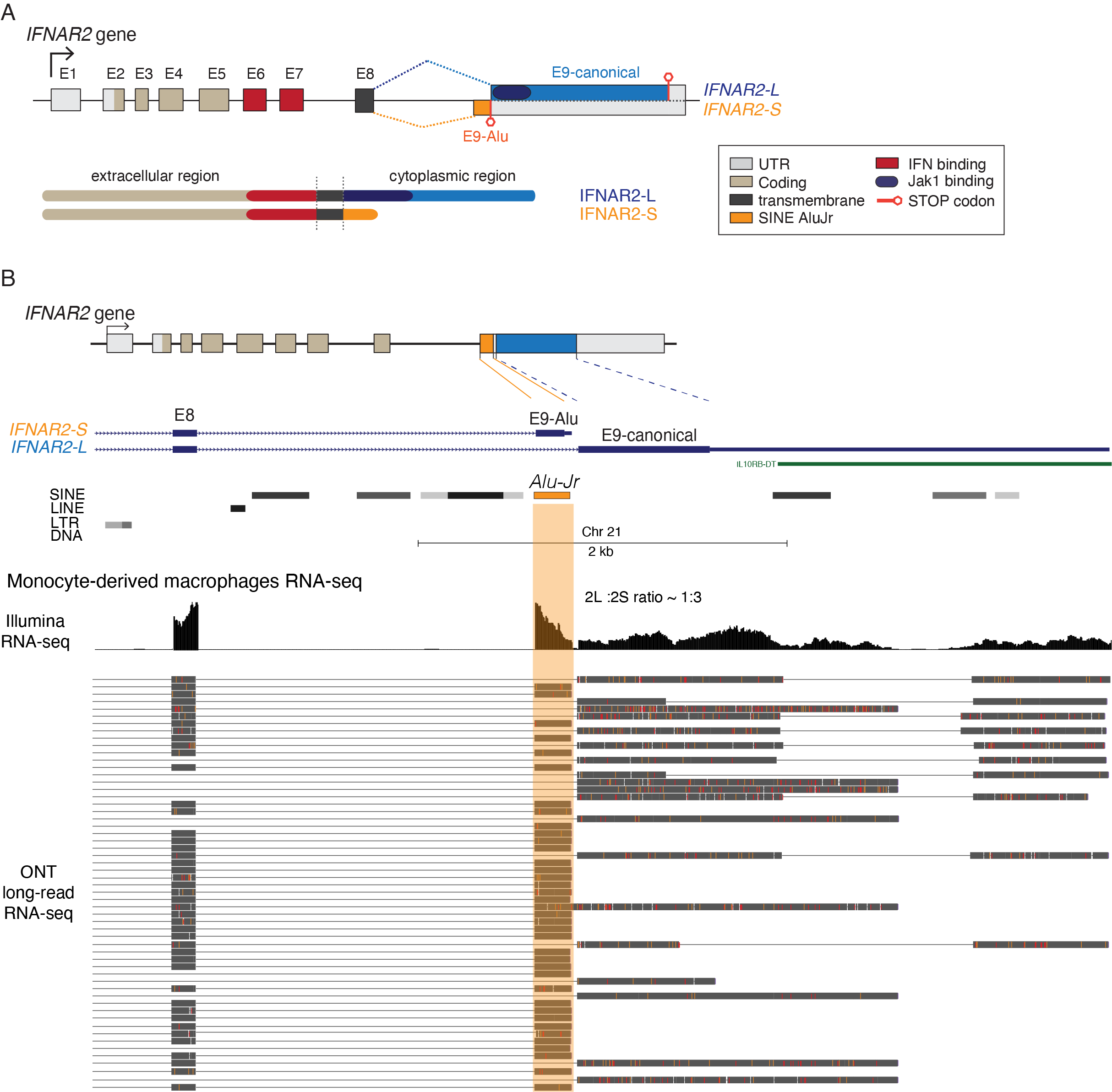

**Fig. S2) Human IFNAR2 is constitutively spliced into two main isoforms.** **A)** Schematic representation of the type-I interferon receptor chain 2 (*IFNAR2*) gene. On top, alternative splicing patterns; at the bottom, schematic of the encoded peptides. **B)** UCSC screenshot shows paired short-read (Illumina) RNA-seq and Oxford Nanopore (ONT) long-read RNA-seq read coverage over exon 8 (E8), exon 9-Alu (E9-Alu) and the canonical full-length exon 9 (E9-canonical) in monocyte-derived macrophages (data from https://genome.ucsc.edu/s/vollmers/IAMA). Both sequencing technologies show that the truncated *IFNAR2-S* isoform is transcribed at higher levels that the full-length *IFNAR2-L* in immune cells.

Supplementary Figure 3

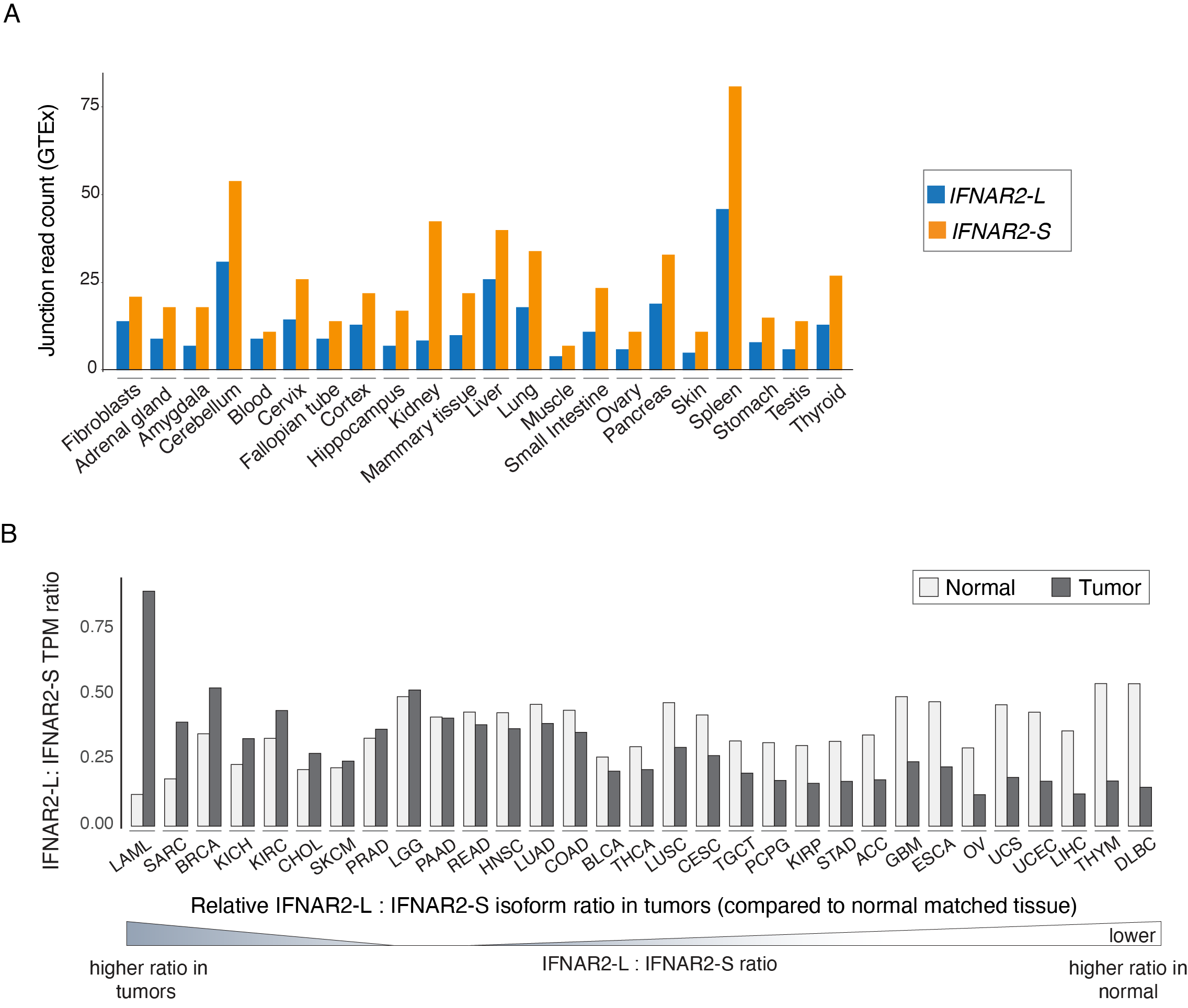

**Fig. S3) Expression of IFNAR2 isoforms in human tissues. A)** Short-read RNA-seq junction read counts comparing *IFNAR2-S* and *IFNAR2-L* expression across normal human tissues. Normalized exon-exon junction read counts from the GTEx analysis V8 (dbGaP Accession phs000424.v8.p2). **B)** Ratios of IFNAR2-L to IFNAR2-S TPM levels across matched healthy (light grey) and tumor (dark grey) samples; isoform count data were accessed through the Gepia2 portal web interface (http://gepia2.cancer-pku.cn). Labels correspond to TCGA study abbreviations for each cancer. TPM = transcripts per million.

Supplementary Figure 4

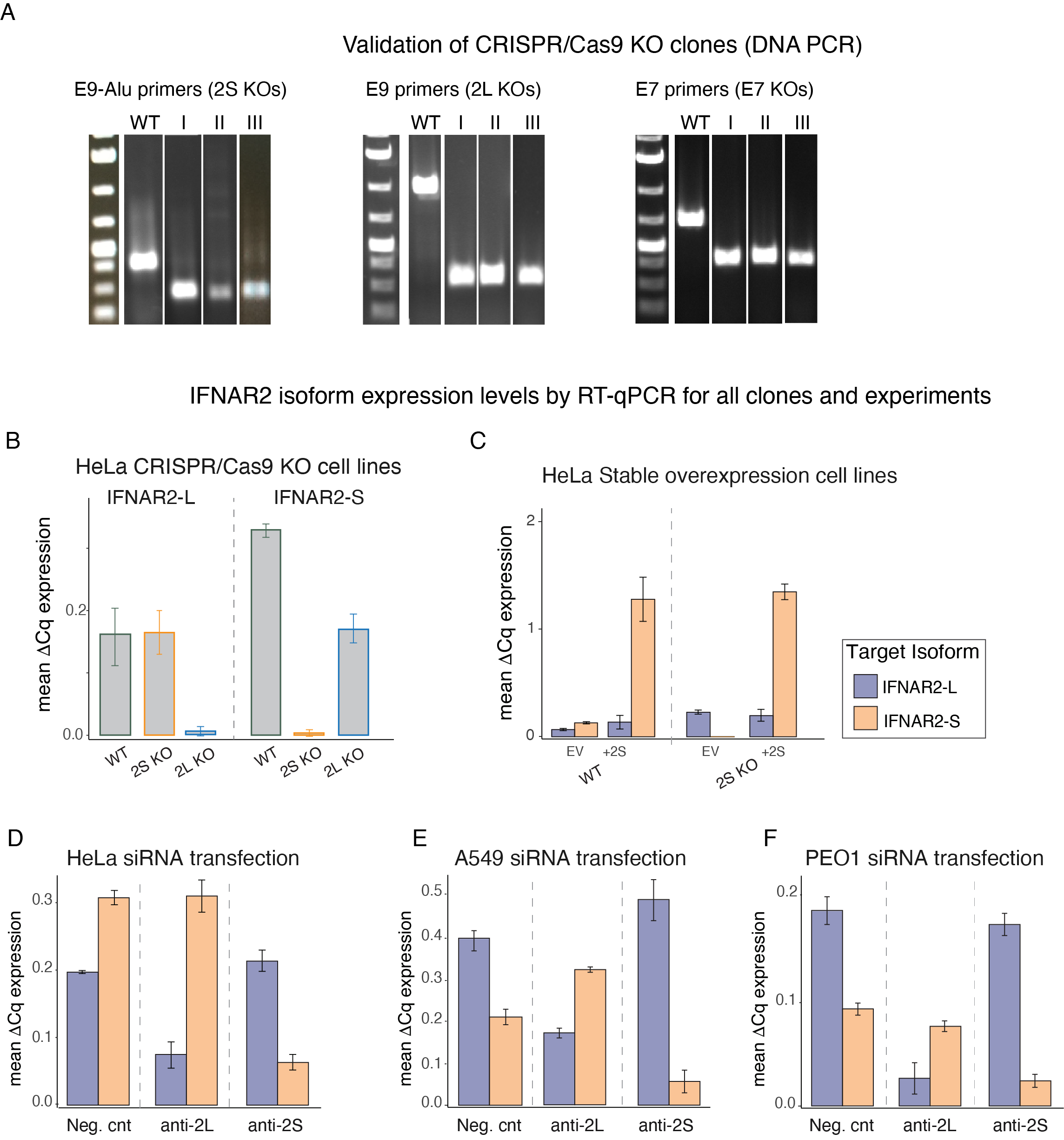

**Fig. S4) Cell line and isoform expression level validation.** **A)** Agarose gel electrophoresis results confirming exon deletion in 3 HeLa homozygous KO clones per genotype. **B-F)** Mean expression levels of the IFNAR2-S (orange) and IFNAR2-L (blue) isoforms across cell lines and experimental protocols. Values represent averaged isoform-specific Cq normalized to the housekeeping gene *CTCF* (ΔCt) as assayed by RT-qPCR for: **B)** wild-type (WT) and knockout HeLa monoclonal cell lines, **C)** wild-type and KO HeLa cells stably transfected with an empty vector (EV) or with the IFNAR2-S isoform (+2S); **D-F)** HeLa cells, A549 and PEO1 cells transfected with either a negative control siRNA (Neg. cnt) or isoform-specific siRNAs (anti-2L and anti-2S).

Supplementary Figure 5

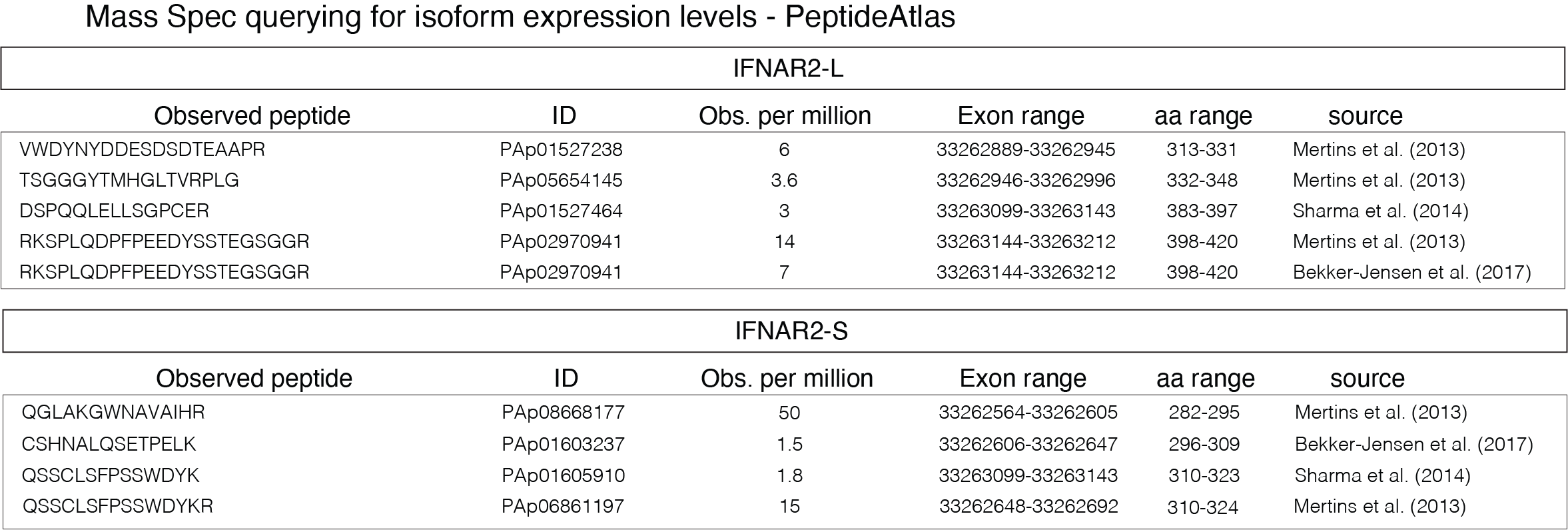

**Fig. S5) Protein expression levels of IFNAR2 isoforms from Mass Spec data.** Support for the translation of both IFNAR2-S and IFNAR2-L into proteins from Mass Spec data (source: The PeptideAtlas project)

Supplementary Figure 6
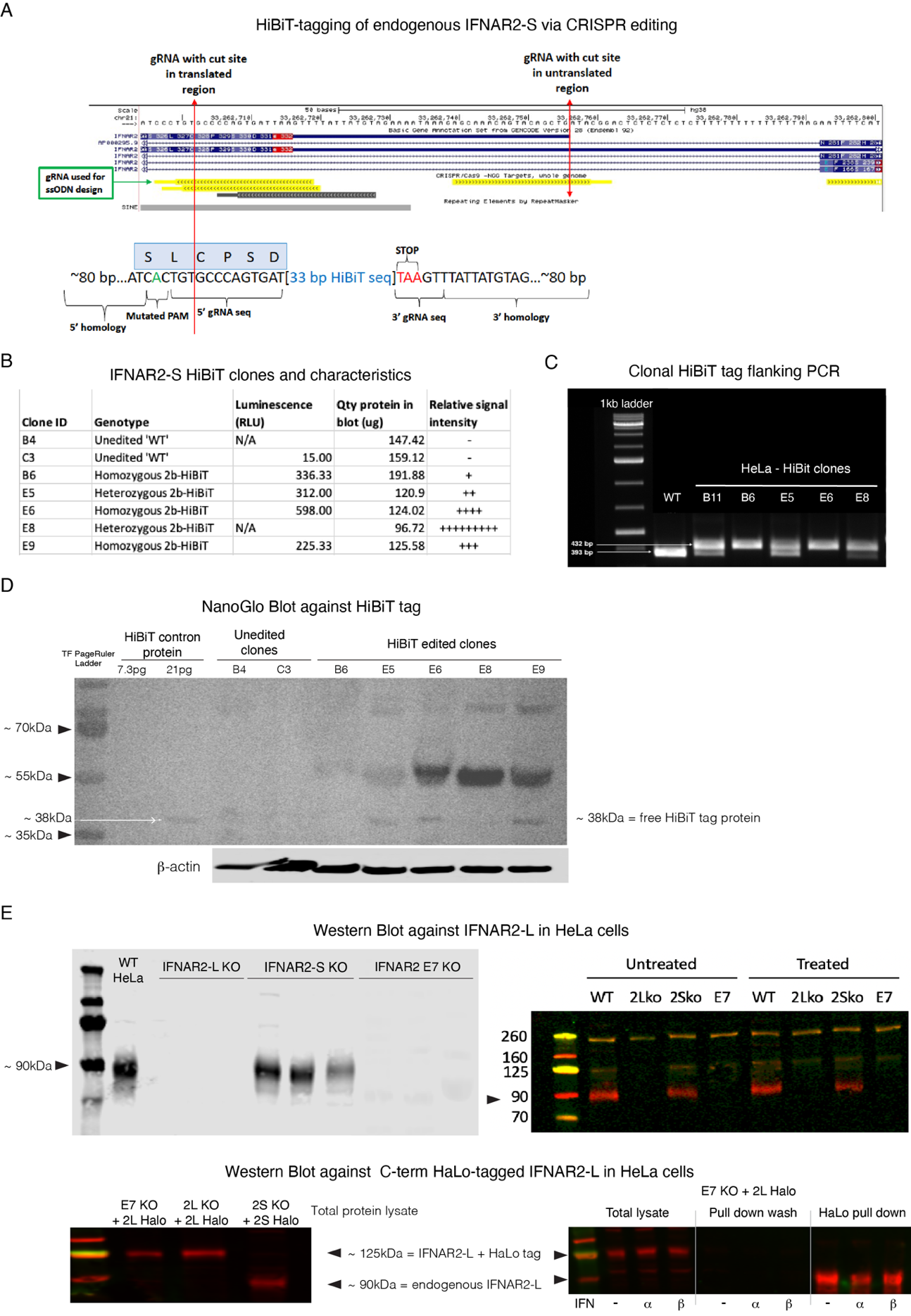

**Fig. S6) Protein expression levels of IFNAR2 isoforms in HeLa cells.** Due to the unavailability of commercial antibodies specific for IFNAR2-S we used CRISPR to add an epitope tag (HiBiT, Promega) upstream of the STOP codon of the IFNAR2-S isoform in wild-type HeLa cells. **A)** Detail of the HiBiT tag editing approach. **B)** Results of the Nano-Glo HiBiT Lytic Detection System used to screen clonally expanded HeLa cells for the integration of the HiBiT tag. **C)** Positive clones from (B) were further expanded and genotyped by PCR and Sanger sequencing. Higher molecular weight bands (432bp) correspond to HiBiT tagged IFNAR2-S isoforms; lower molecular weight bands (393 bp) correspond to the amplicon of wild-type IFNAR2-S isoforms. Clones with both bands are heterozygous for IFNAR2-S (B11, E5, E8) and were selected because of high HiBiT tag detection levels in (B). **D)** Validation by HiBiT blotting of IFNAR2-S translation as protein. The ~38kDa band corresponds to the HiBiT-LgBiT protein complex. **E)** **Top**: Western blots show expression of IFNAR2-L in wild-type (WT) and IFNAR2-S KO (2S KO) HeLa cells, but not in IFNAR2-L KO (2L KO) and IFNAR2 KO (E7 KO) cells. **Bottom**: Western blots show expression of IFNAR2-L in IFNAR2 KO and IFNAR2-L KO cells stably overexpressing a C-terminal Halo-tagged version of IFNAR2-L. In the total protein lysate fraction (left) the HaloTag IFNAR2-L isoform can be identified as a ~125kDa protein (IFNAR2-L= ~90kDa; HaloTag = ~33kDa). IFNAR2-S KO cells were used as positive control. On the right, IFNAR2-L was correctly retrieved in the pull-down fraction as a 90kDa protein upon cleavage of the HaloTag.

Supplementary Figure 7

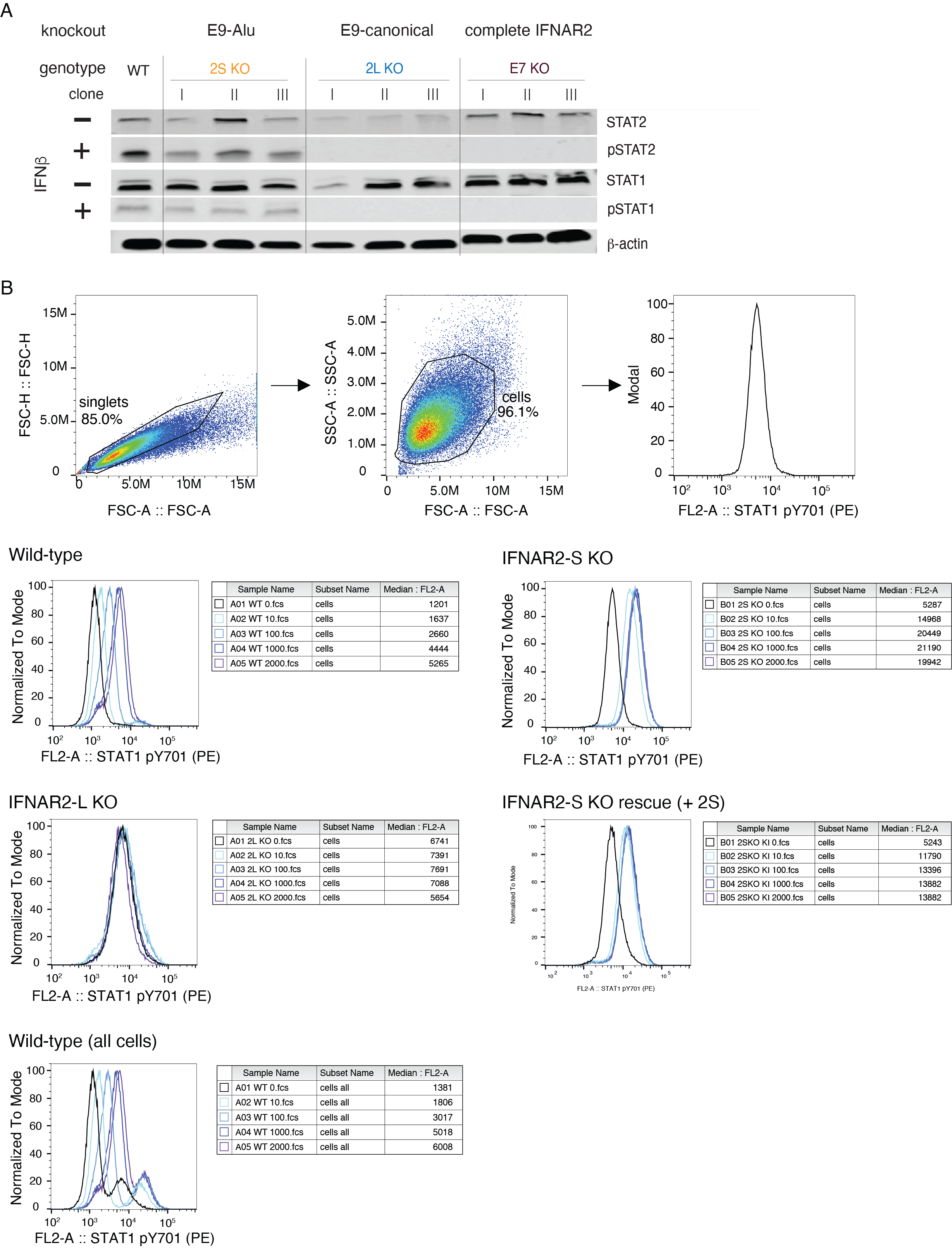

**Fig. S7) IFNAR2-mediated phosphorylation of STAT1 and STAT2. A)** Total and phosphorylated STAT1 (pSTAT1) and STAT2 (pSTAT2) levels were assessed by western blot in wild-type and knockout HeLa cell lines upon 30min of IFNβ stimulation (10U/ml). For each KO cell line, 3 different validated clones were screened. Our data consistently show IFNAR2-L dependent STAT1/2 phosphorylation, as illustrated by lack of pSTAT1 and pSTAT2 detection in IFNAR2-L and IFNAR2 KO clones. **B)** STAT1 phosphorylation was assessed upon treatment for 30min with increasing doses of IFNβ (0U, 10U/ml, 100U/ml, 1000U/ml and 2000U/ml) by flow cytometry using a primary antibody specific for intracellular pSTAT1 staining and detection (pY701-PE). On top, a description of how the events to analyze were collected: singlets and then cells. Cells were analyzed for median fluorescence intensity (MFI) in the PE channel. Untreated (0U/ml) wild-type cells were used as reference to define the range comprising the increase of pSTAT1 signal. Panels at the bottom show the signal acquired for each cell line at the different treatment conditions. Wild-type cells show increasing response to IFNβ (shift toward the right in median peak) at increasing concentration of IFNβ. In contrast, IFNAR2-L KO cells showed no increase in pSTAT1 signal, whereas IFNAR2-S KO cells not only show higher levels of pSTAT1 than wild-type cells at the lowest dose of IFNβ, but also reached signal saturation at 100U/ml of IFNβ. At the very bottom, we show pSTAT1 signal from all cells, meaning to include a subpopulation among gfated singlets that was present only for this cell type. This expanded population shows similar results to only gating on the major population, and this data was not used.

Supplementary Figure 8

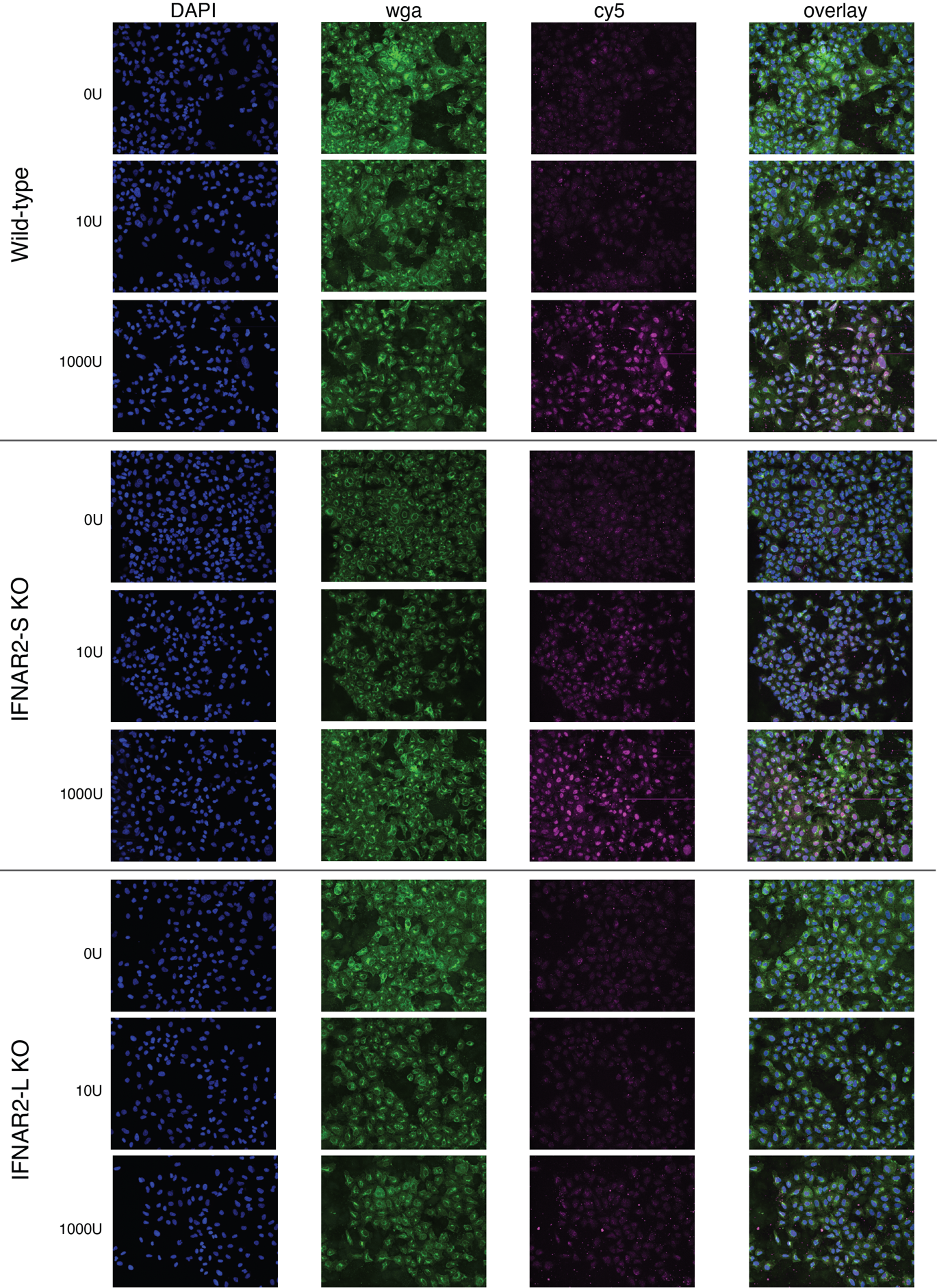

**Fig. S8) Immunofluorescence detection of phosphorylated STAT2.** Phosphorylated STAT2 (pSTAT2) signal was collected from wild-type and knockout A549 cell line treated with increasing concentration of IFNβ (0U/ml, 10U/ml and 1000U/ml) for 5min. In agreement with previous analyses of STAT phosphorylation, we detected higher levels of pSTAT2 in the nucleus of IFNAR2-S KO cells compared to wild-type cells at lower concentrations of IFNβ treatment (10U/ml). No signal was detected in IFNAR2-L KO cells.

Scale: 368 X 460 μm. Blue channel = DAPI (nuclear stain). Green channel = Wheat Germ Agglutinin (wga) stain in the GFP channel (membrane stain). Pink channel = Cy5 (pSTAT2 stain).

Supplementary Figure 9

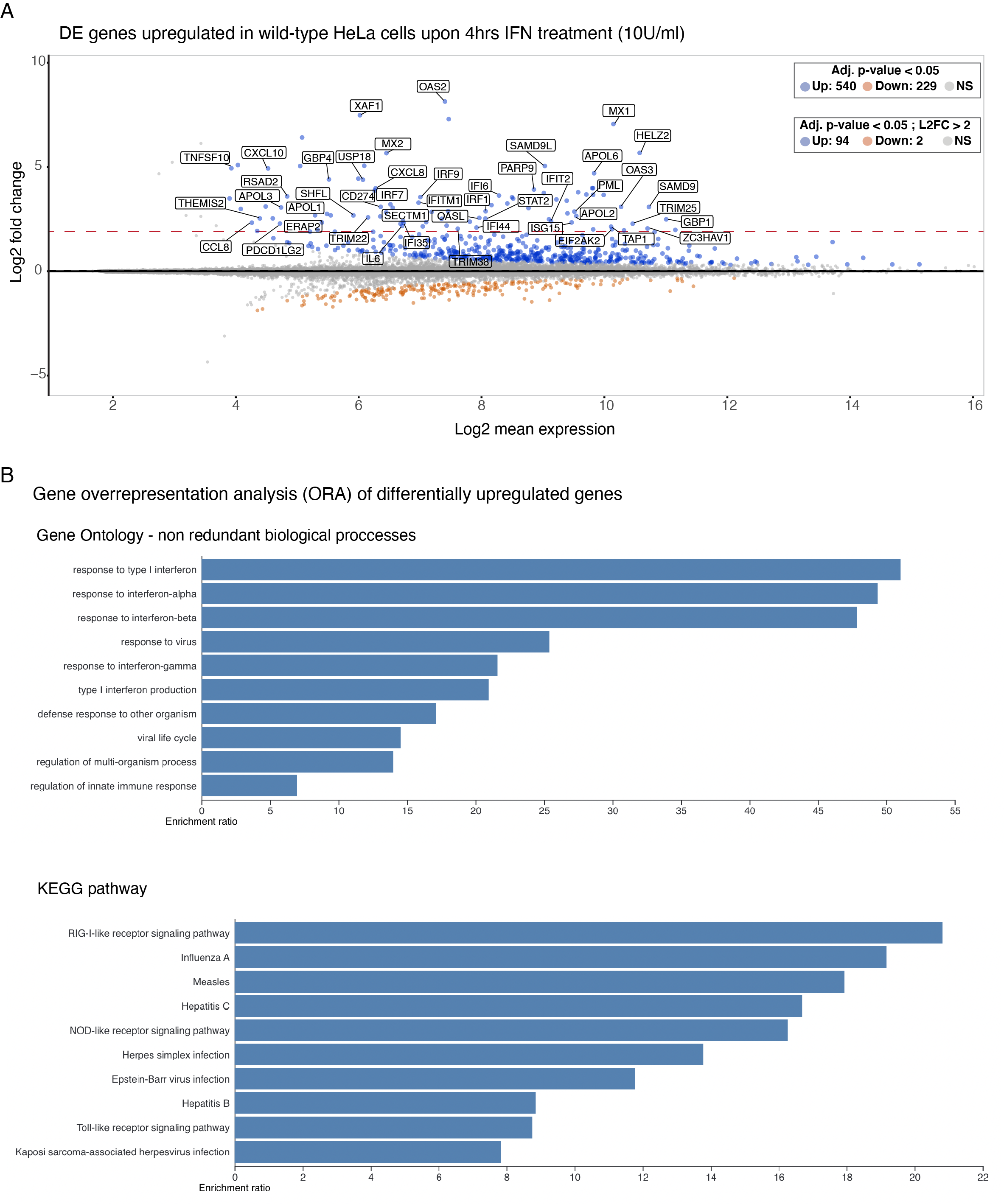

**Fig. S9) Differential gene expression analysis for HeLa wild-type cells. A**) MAplot shows differentially upregulated genes (adj. p-value < 0.05 and log2 fold change >2) in IFNβ treated vs. untreated HeLa wild-type cells. **B**) Gene overrepresentation analyses of differentially upregulated genes in IFNβ treated cells for non-redundant biological processes and Kegg pathways. Only significant enrichment is reported (false discovery rate < 0.05). Overrepresentation analyses were performed using the WebGestalt portal (http://www.webgestalt.org/)

Supplementary Figure 10

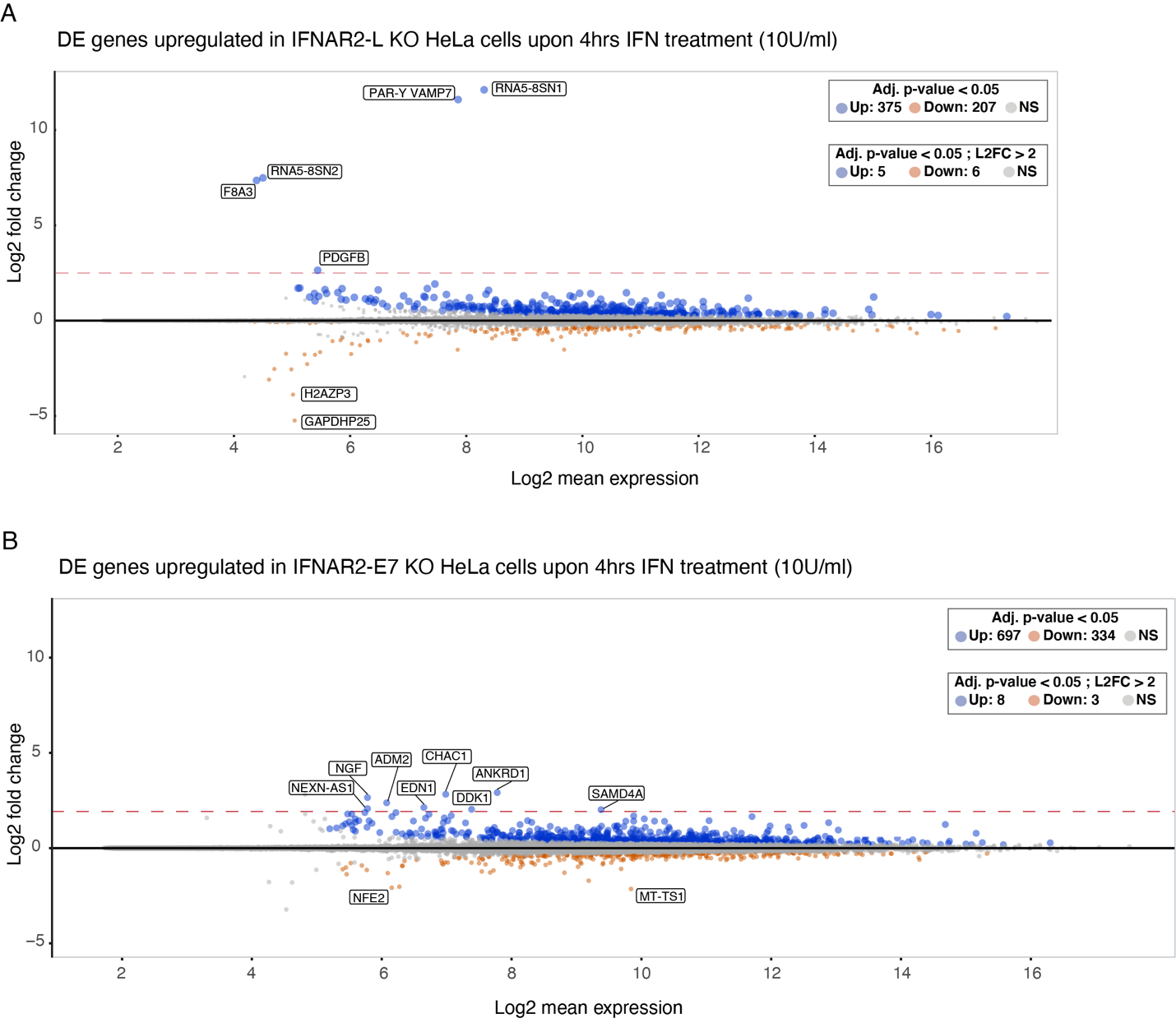

**Fig. S10) Differential gene expression analysis for IFNAR2-L KO and IFNAR2 KO HeLa cells. A)** MAplot shows differentially upregulated genes (adj. p-value < 0.05 and log2 fold change >2) in IFNβ treated vs. untreated IFNAR2-L KO cells. **B)** MAplot shows differentially upregulated genes (adj. p-value < 0.05 and log2 fold change >2) in IFNβ treated vs. untreated IFNAR2 KO cells.

No gene ontology enrichment terms were identified.

Supplementary Figure 11

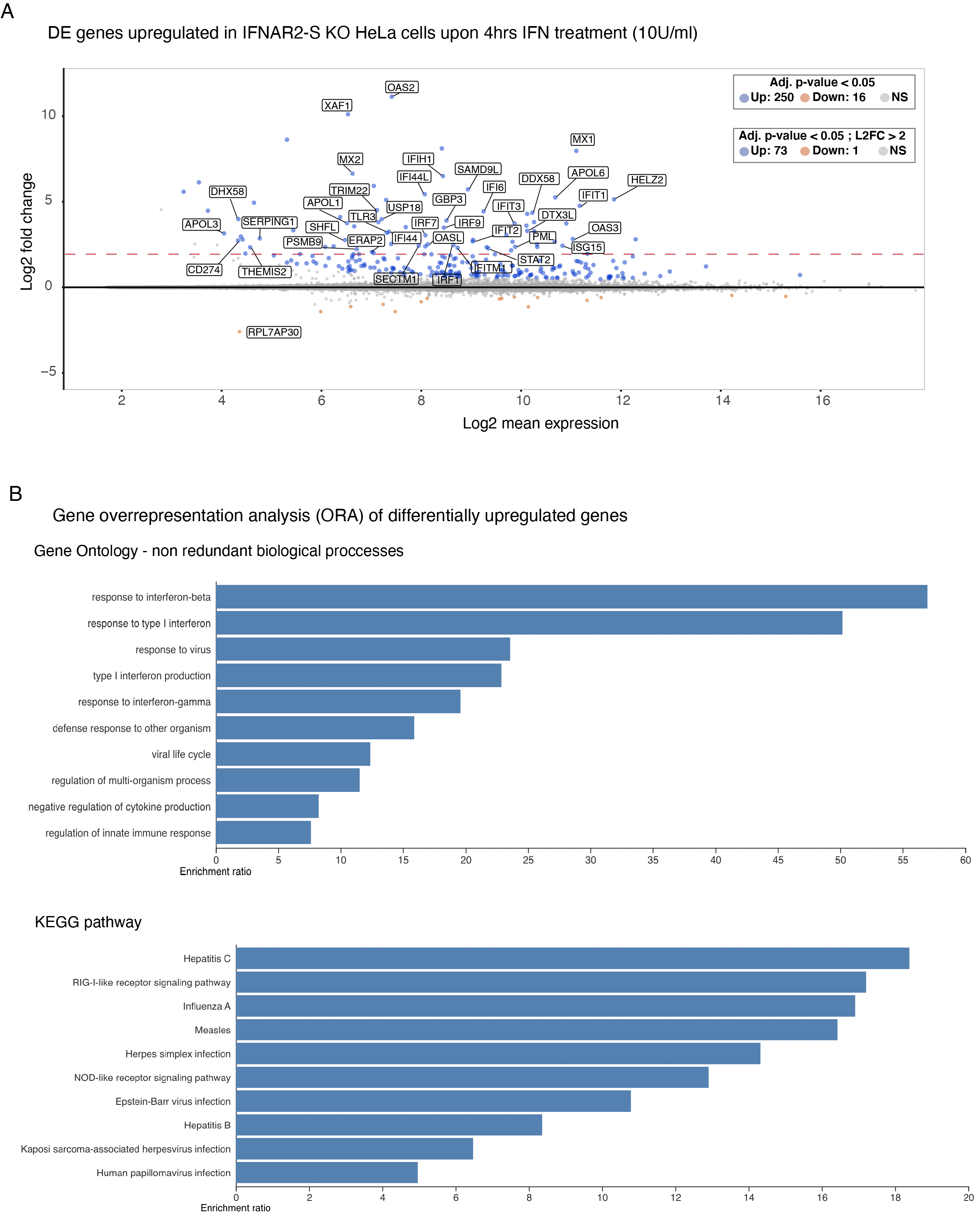

**Fig. S11) Differential gene expression analysis for IFNAR2-S KO HeLa cells. A)** MAplot shows differentially upregulated genes (adj. p-value < 0.05 and log2 fold change >2) in IFNβ treated vs. untreated IFNAR2-S KO HeLa cells. **B)** Gene overrepresentation analyses of differentially upregulated genes in IFNβ treated cells for non-redundant biological processes and Kegg pathways. Only significant enrichment is reported (false discovery rate < 0.05).

Overrepresentation analyses were performed using the WebGestalt portal (http://www.webgestalt.org/)

Supplementary Figure 12

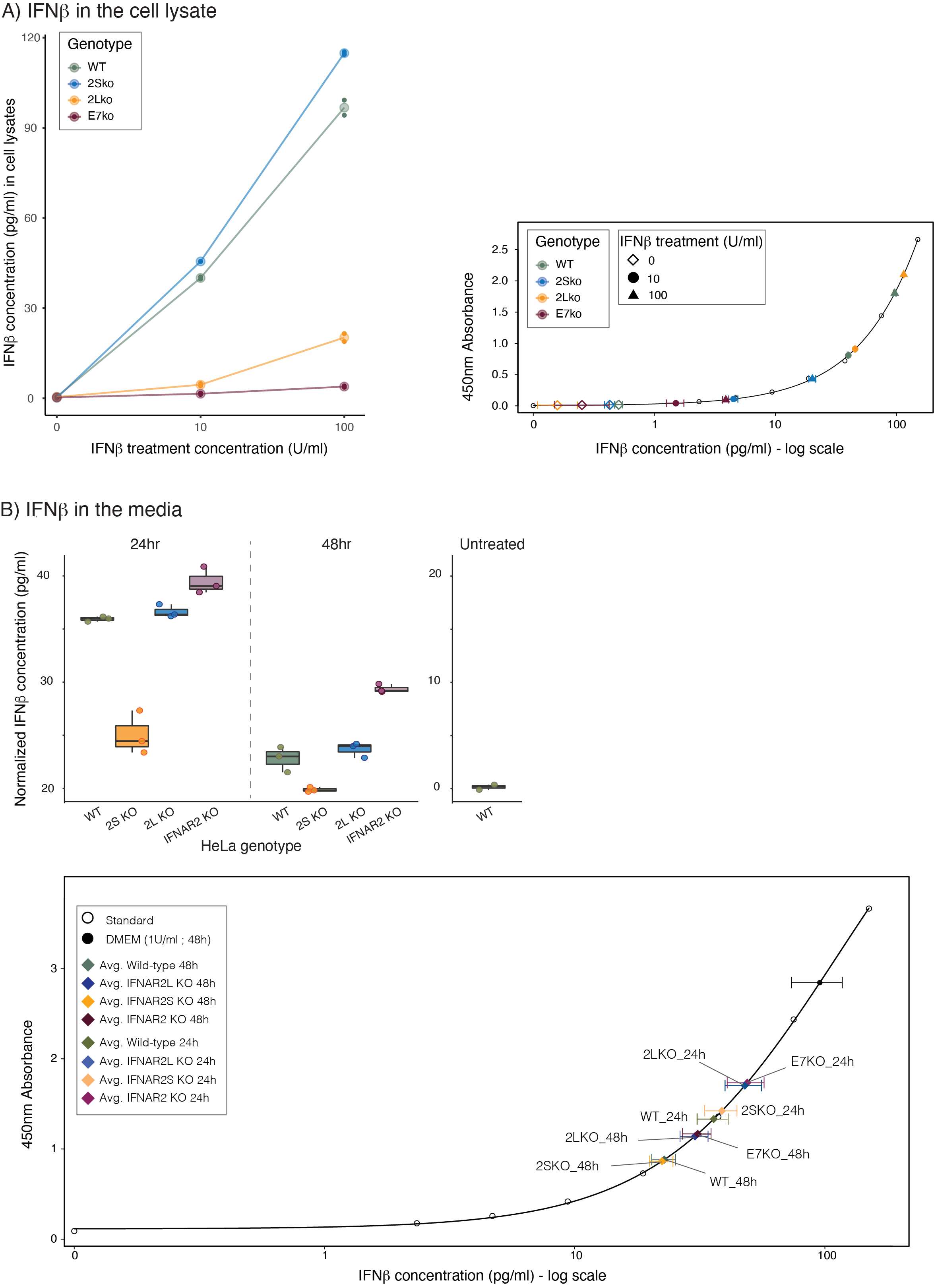

**Fig. S12) Cellular IFNβ intake as measured by ELISA. A)** Histogram shows the concentration of IFNb internalized by wild-type and mutant HeLa cell lines upon 30min treatment with increasing doses of IFNβ. After treatment, we performed total protein extraction and equal amounts of protein were used to perform the ELISA assay. Absorbance values from triplicates were converted into IFNβ concentration using the standard curve on the right. **B**) ELISA measurements of the normalized concentration of recombinant IFNβ remaining in media after 24hrs or 48hrs treatment with 1U/ml (~60pg/ml) of IFNβ. Right panel shows that wild-type HeLa cells (WT) do not secrete IFNβ. At the bottom is reported the standard curve used to convert absorbance to IFNβ concentration.

Supplementary Figure 13

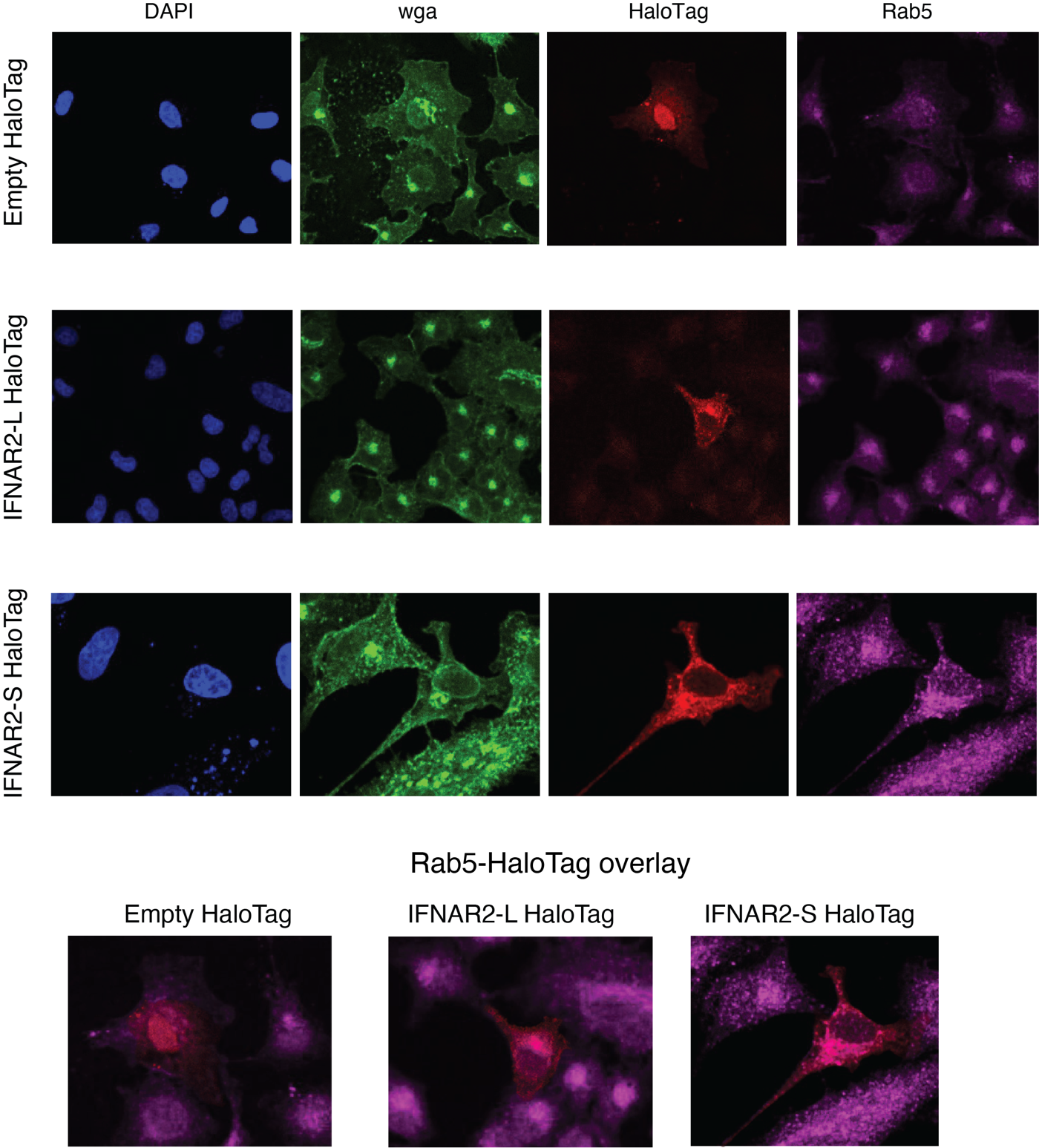

**Fig. S13) IFNAR2 isoforms co-localize with endosomal proteins (Rab5) upon IFNβ stimulation**. Rab5 and HaloTag immunofluorescent (IF) signal was collected from wild-type A549 cell line transiently expressing an empty HaloTag vector (Empty HaloTag; top), IFNAR2-L HaloTag (C-term; middle) or IFNAR2-S HaloTag (C-term; bottom). Cells were treated for 5min with 10U/ml of IFNβ before imaging. Scale: 184 um x 230 μm.

Blue channel = DAPI (nuclear stain). Green channel = Wheat Germ Agglutinin (wga) stain in the GFP channel (membrane stain). Pink channel = Rab5. Red channel = JFX549 permeable HaloTag ligand.

Supplementary Figure 14

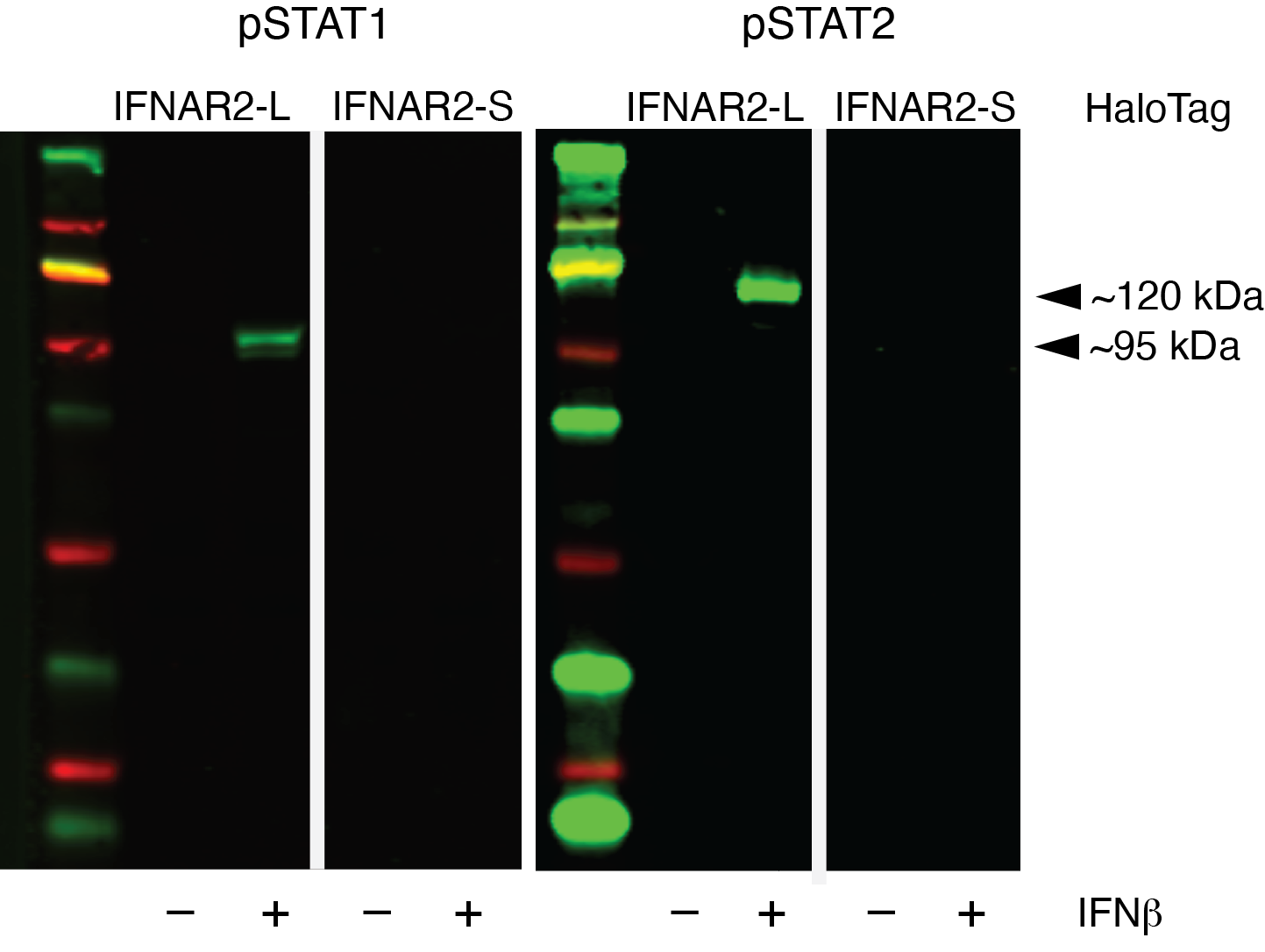

**Fig. S14) The C-terminal HaloTag does not interfere with the activation of type I IFN signaling.** Western Blot targeting phosphorylated STAT1 (left) and STAT2 (right) in untreated (IFNβ -) and IFNβ treated cells (10U/ml for 30 min; IFNβ +). To confirm that the signal detected was coming specifically from the HaloTag isoform, we transfected IFNAR2 KO cells with either a IFNAR2-L C-terminal HaloTag (HaloTag: IFNAR2-L) or a IFNAR2-S C-terminal HaloTag (HaloTag: IFNAR2-S). We have previously shown (Fig. S7A) that IFNAR2 KO cells do not phosphorylate STAT1/2 upon IFN treatment.

Supplementary Figure 15

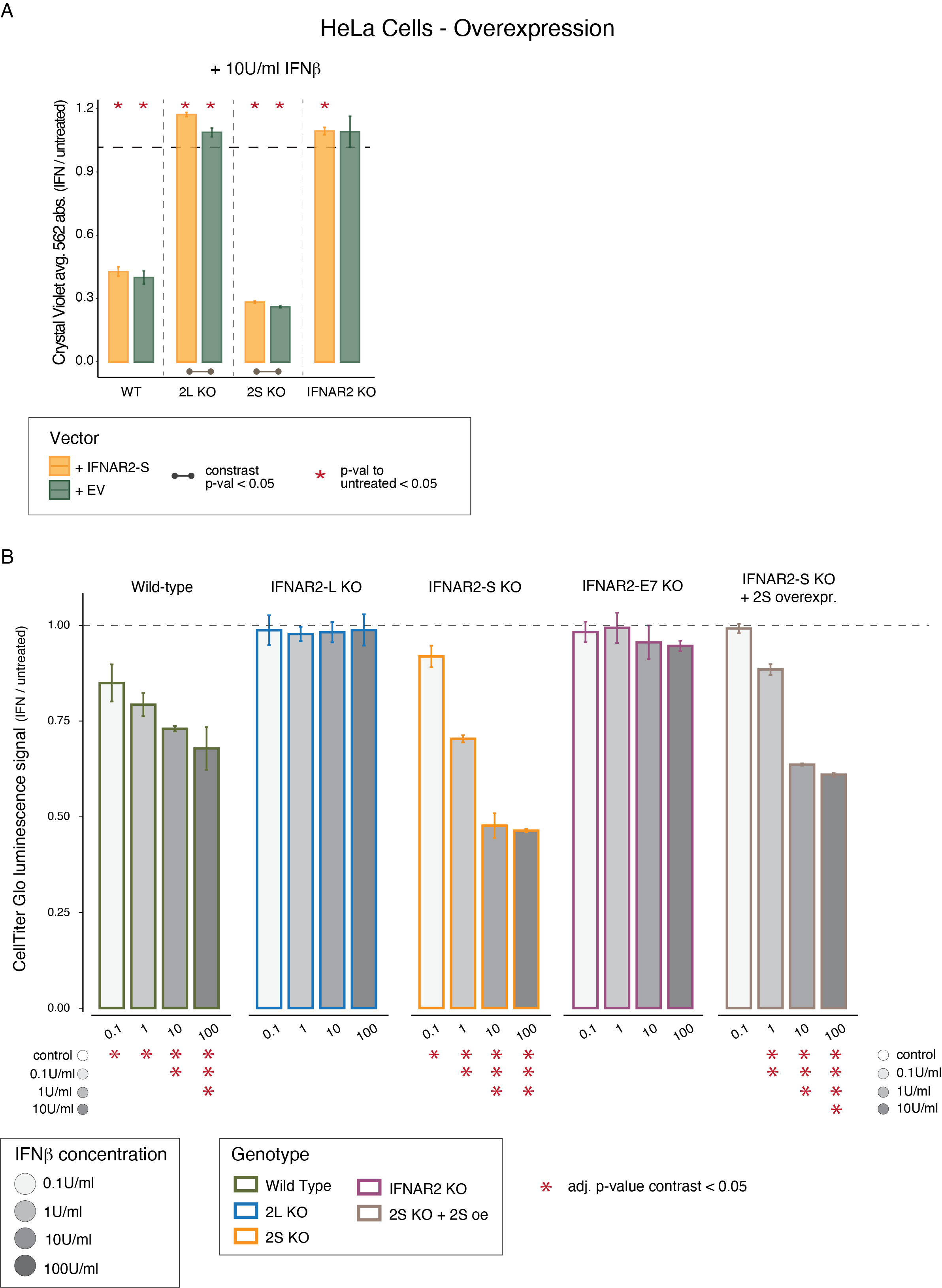

**Fig. S15) The IFNAR2-L : IFNAR2-S ratio affects cellular sensitivity to cytotoxic effects of IFNβ treatment.** **A)** Histogram shows changes in cell viability and proliferation as assessed by crystal violet staining in wild-type (WT) and knockout HeLa cells transfected with an empty vector (EV, in green) or a vector for the stable expression of IFNAR2-S (+IFNAR2-S, in yellow) treated for 4 day with IFNβ compared to untreated cells. Asterisks (*) on top of each bar represent a significant difference between untreated and treated cells; Bars at the bottom represent a significant difference in contrast between EV and +2S vector based on an ANOVA *emmeans* contrast statistical test. **B)** Histograms show the loss of cell viability upon IFN treatment compared to untreated cells as measured by CellTiter Glo assay of cell viability. Cells were left for 4 days at increasing doses of IFNβ (0.1U/ml to 100U/ml) or in complete media without IFN (untreated) before reading luminescence. Asterisks (*) at the bottom represent a significant adj. p-value in within-genotype contrasts between treatment conditions based on an ANOVA *emmeans* contrast statistical test.

Supplementary Figure 16

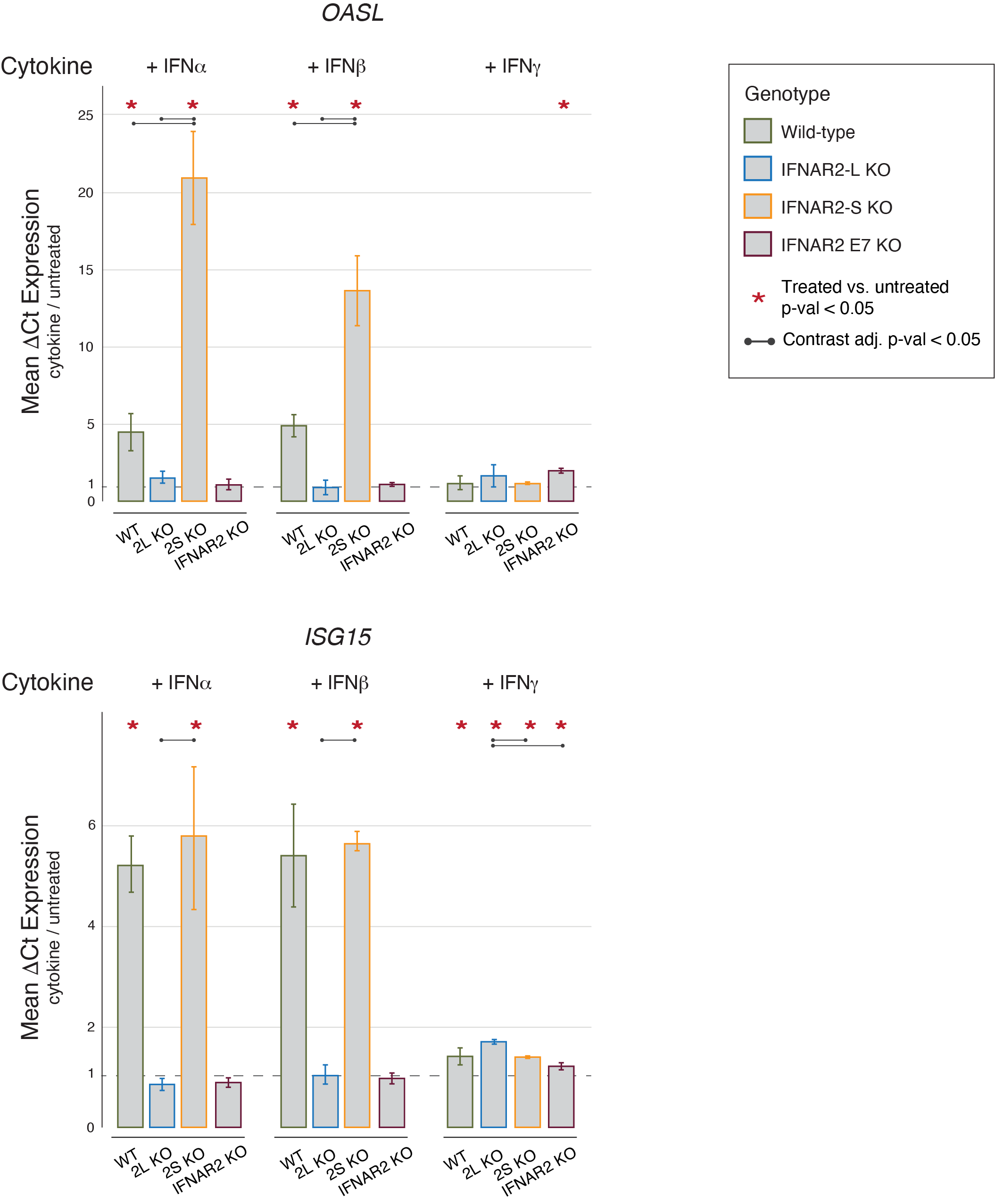

**Fig. S16) ISG expression levels in wild-type and KO HeLa cell lines treated with different cytokines.** Histograms show the mean expression levels (ΔCt) of *OASL* (top) and *ISG15* (bottom) between treated and untreated cells as measured by RT-qPCR. ΔCt were calculated by normalizing the Ct of the target gene by the Ct of the housekeeping CTCF gene. Cells were treated for 4 hours with 100U/ml of different cytokines (type I: IFNα and IFNβ. type II:

IFNγ) or equal volume of DPBS (untreated control) before RNA extraction.

Supplementary Figure 17

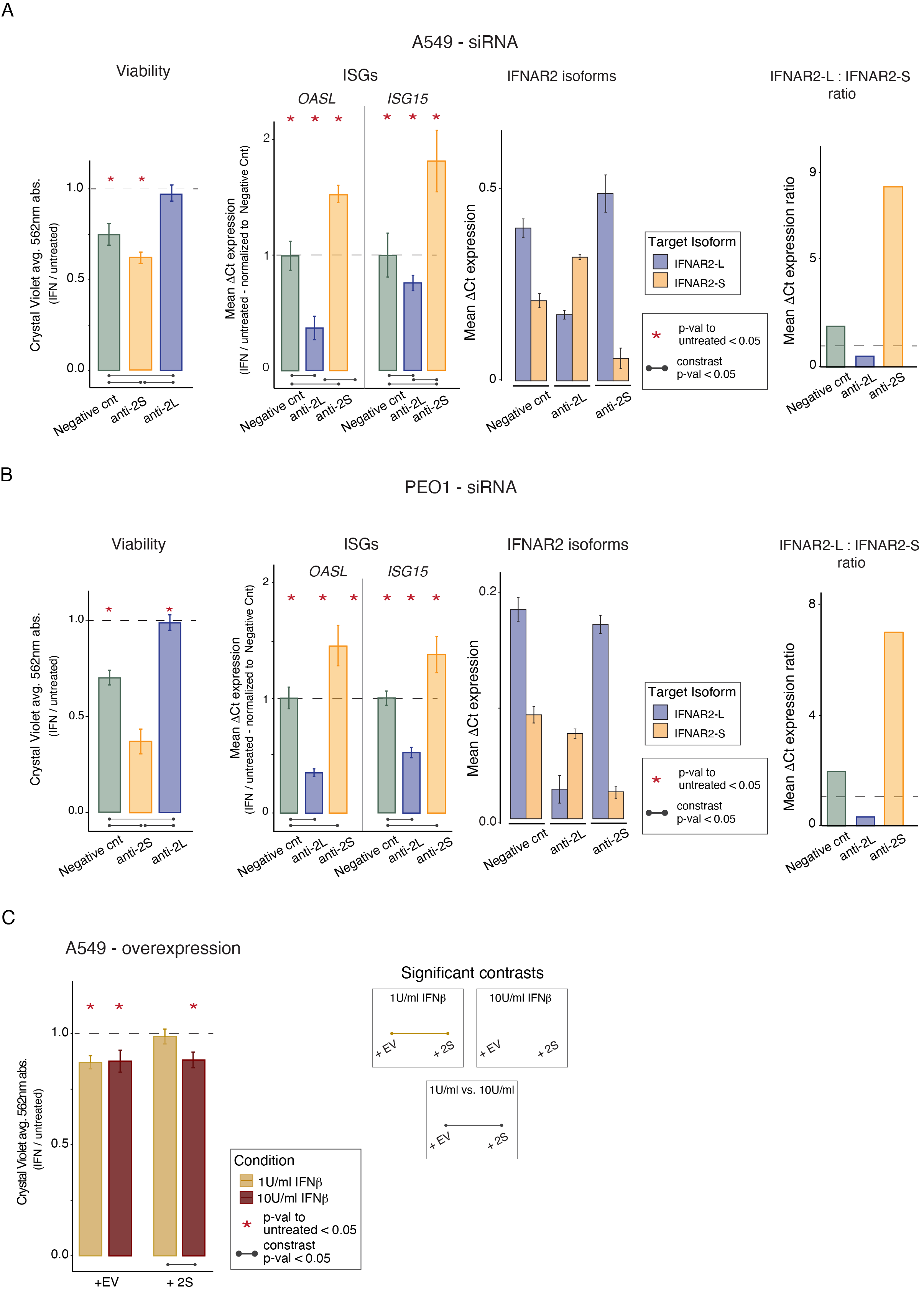

**Fig. S17)** **Effect of IFNβ treatment in additional cell lines – A549 and PEO1. A-B)** Changes in viability and expression levels of the canonical ISGs *OASL* and *ISG15* (mean ΔCt) between IFNβ-treated and untreated cells. ΔCt values were calculated by normalizing the Ct of the target gene to the Ct of *CTCF* (housekeeping gene) for the same sample and replicate. A549 (**A**) and PEO1 cells (**B**) were transfected with a negative control and isoform-specific siRNAs (anti-2L and anti-2S) 48hrs before being treated with IFNβ for 4 days (viability) or 4hrs (RT-qPCR of ISGs) or equal volume of DPBS (untreated control). Effectiveness and stability of the siRNA transfection was verified by RT-qPCR for IFNAR2-L (blue) and IFNAR2-S (orange), and reported as mean isoform expression or as relative expression ratio (ΔCt IFNAR2-L / ΔCt IFNAR2-S). **C**) Results of cell viability assay as measured by crystal violet absorbance in A549 cells overexpressing either an empty vector (+EV) or IFNAR2-S (+2S). Cells were treated for 4 days with either 1U/ml (gold) or 10U/ml (red) of IFNβ, and their viability and proliferation was compared to that of matched untreated cells. Asterisks (*) indicate a significant difference in treated cells compared to untreated controls. Horizontal bars indicate a significance difference between contrast groups based on an ANOVA *emmeans* pairwise contrast statistical test.

Supplementary Figure 18

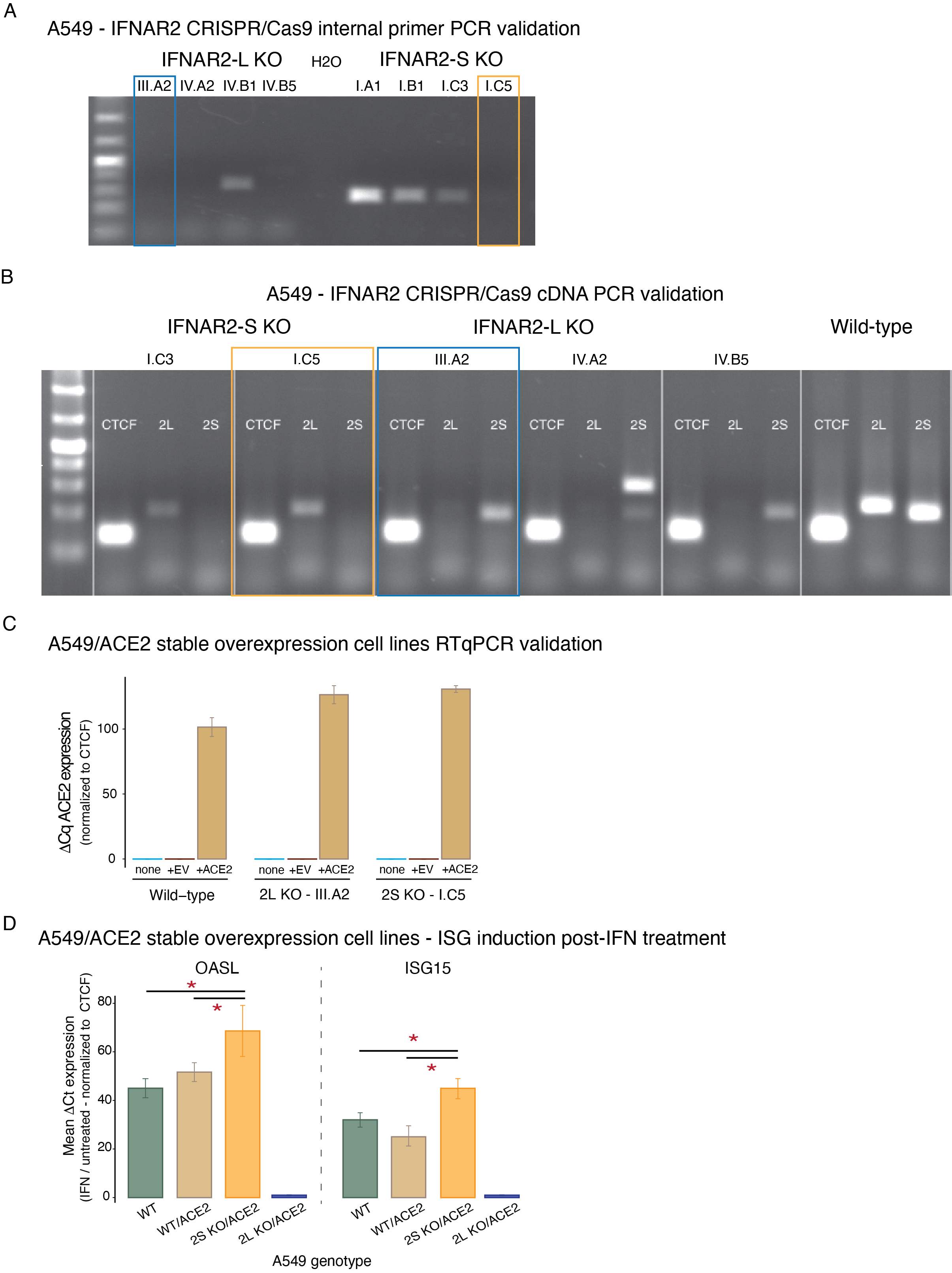

**Fig. S18) Validation of IFNAR2-L and IFNAR2-S knockout A549 cell lines stably expressing ACE2.** **A)** Agarose gel electrophoresis using internal primers confirms in at least one clone a homozygous target-exon deletion (KO) per genotype (highlighted in blue -IFNAR2-L KO- and yellow -IFNAR2-S KO-). **B)** Agarose gel electrophoresis on cDNA of selected IFNAR2-L and IFNAR2-S clones further confirms that deletion of one exon did not affect the transcription of the alternative isoform. **C)** RT-qPCR was used to verify expression of ACE2 in wild-type and knockout A549 cells transfected with a piggyBac vector for the stable genomic integration of ACE2. **D)** Histograms show mean expression levels of the canonical ISGs *OASL* and *ISG15* (ΔCt) upon 4hrs of IFNβ treatment (10U/ml) in A549/ACE2 clones normalized to ΔCt values of untreated clones. ΔCt values were calculated by normalizing the Ct of the target gene to the Ct of *CTCF* (housekeeping gene) for the same sample and replicate. This allowed us to verify that effects of IFNβ treatment were consistent between cell lines (HeLa, A549 and A549 transfected with ACE2). Asterisks (*) indicate a significant difference between contrast groups (indicated by the horizontal bars) based on an ANOVA *emmeans* pairwise contrast statistical test.

Supplementary Figure 19

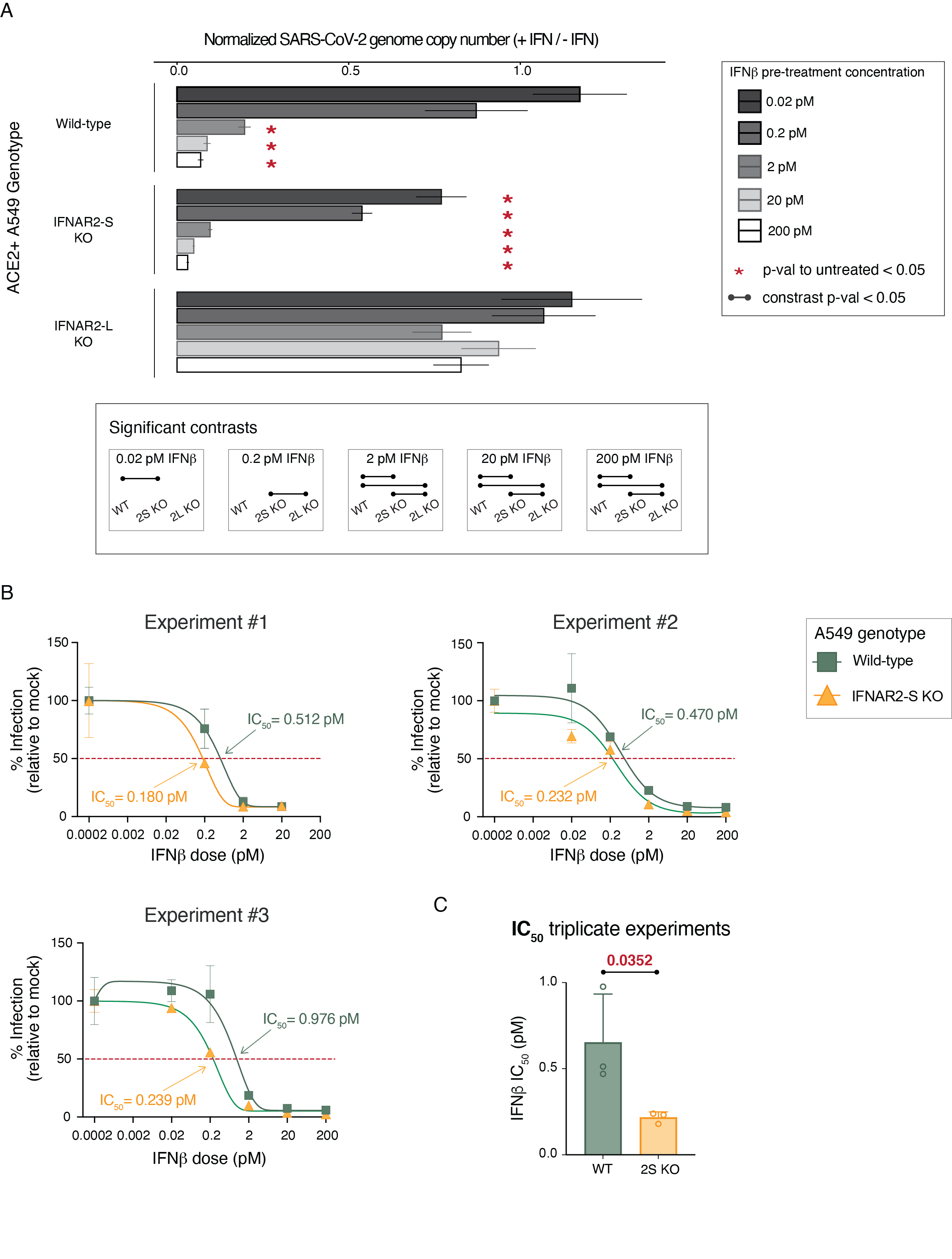

**Supplementary Fig. 19) Protective effect of IFNAR2-L against SARS-Cov-2 infection. A)** Histograms show infection levels of the SARS-Cov-2 virus as a fraction of viral load in IFNβ pre-stimulated cells normalized to viral load in unstimulated cells. Viral load was calculated using a standard curve of SARS-Cov-2 genome copies by RT-qPCR. Even with the lowest doses of IFNβ, cells expressing only IFNAR2-L (IFNAR2-S KO) were able to counter viral infection (normalized viral load <1), whereas wild-type cells show a significant decrease in viral replication levels only when pre-stimulated with >2pM of IFNβ. As expected, cells lacking IFNAR2-L (KO) showed no response to IFNβ pre-treatment, and similar levels of infection in unstimulated and pre-stimulated cells. **B-C)** For each of the three independent viral infection experiments, curves show the % of viral infection (relative to unstimulated mock controls) when cells were pre-stimulated with increasing concentrations of IFNβ in wild-type and IFNAR2-S KO cells. Curves were used to determine the IFNβ IC_50_ (B) for each experiment, and then combined to test whether there is a significant decrease in IFNβ IC_50_ in IFNAR2-S KO cells compared to wild-type cells (C). IFNβ IC_50_ = concentration of IFNβ (pM) required to lower viral genome load (% of infection) by 50% compared to unstimulated cells (mock).

Supplementary Figure 20

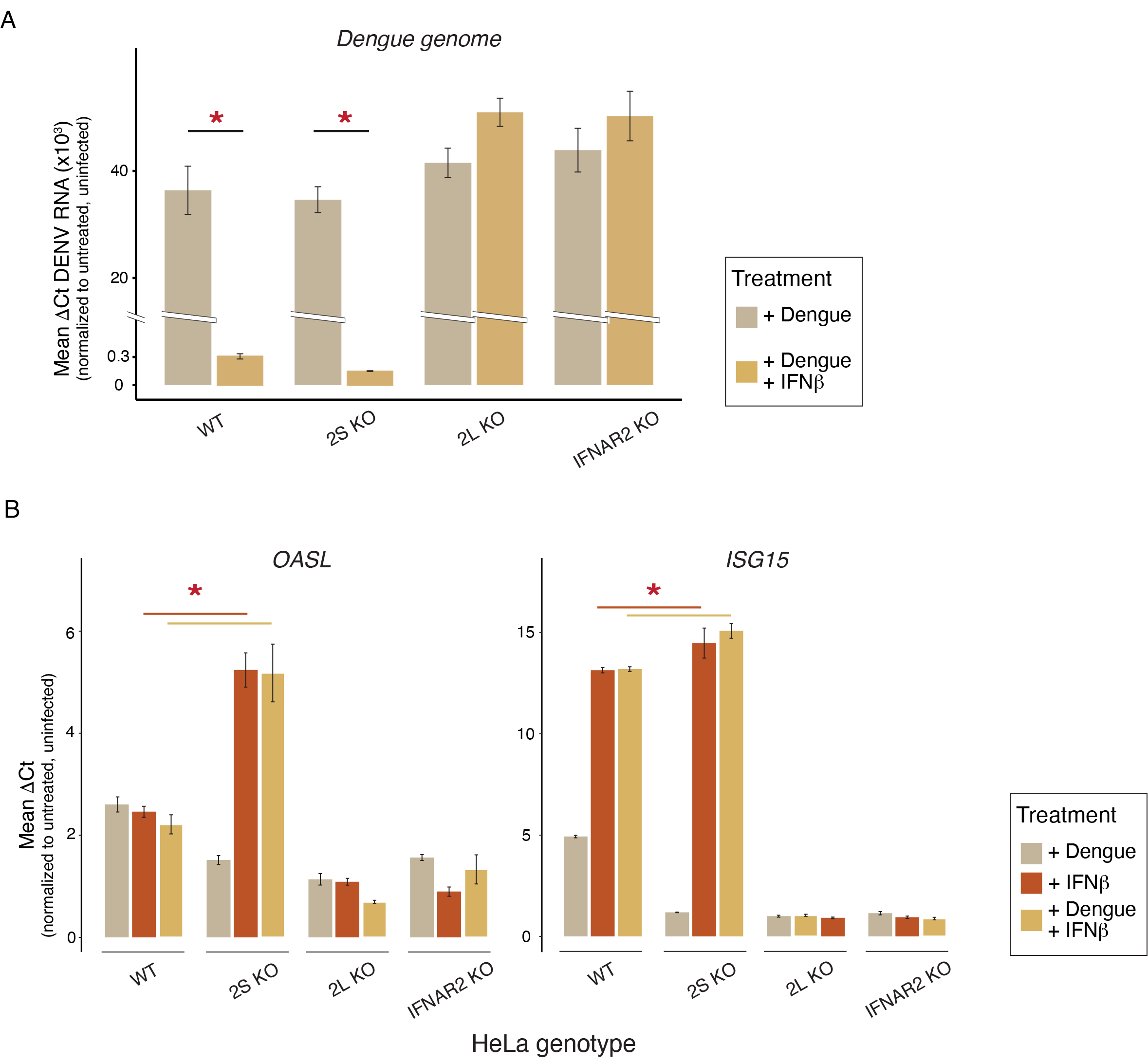

**Fig. S20)** **Protective effect of IFNAR2-L against DENV-2 infection. A)** RT-qPCR measurements of Dengue RNA ΔCt, which correlates with viral genome copies, in lysates of IFNβ pre-stimulated HeLa cells normalized to Dengue RNA ΔCt in unstimulated cells. Whereas no effect of IFNβ pre-stimulation was detected in IFNAR2-L KO and IFNAR2 KO cells, both wild-type and IFNAR2-S cells show significantly lower levels of infection upon IFN pre-stimulation. **B)** From the same samples and experiments, expression levels of canonical ISGs (*OASL* and *ISG15*) were compared between infected cells (+Dengue), pre-stimulated cells (+IFNβ) and pre-stimulated infected cells (+Dengue + IFNβ). Cells were pre-stimulated with 100U/ml IFNβ for 24hr before being infected with DNEV-2 virus (MOI = 1); ΔCt were calculated by normalizing target loci Ct to *CTCF* (housekeeping gene) Ct for the same sample and replicate. Asterisks (*) indicate a significant difference between groups (horizontal bars) according to an ANOVA *emmeans* pairwise contrast statistical test.

Supplementary Figure 21

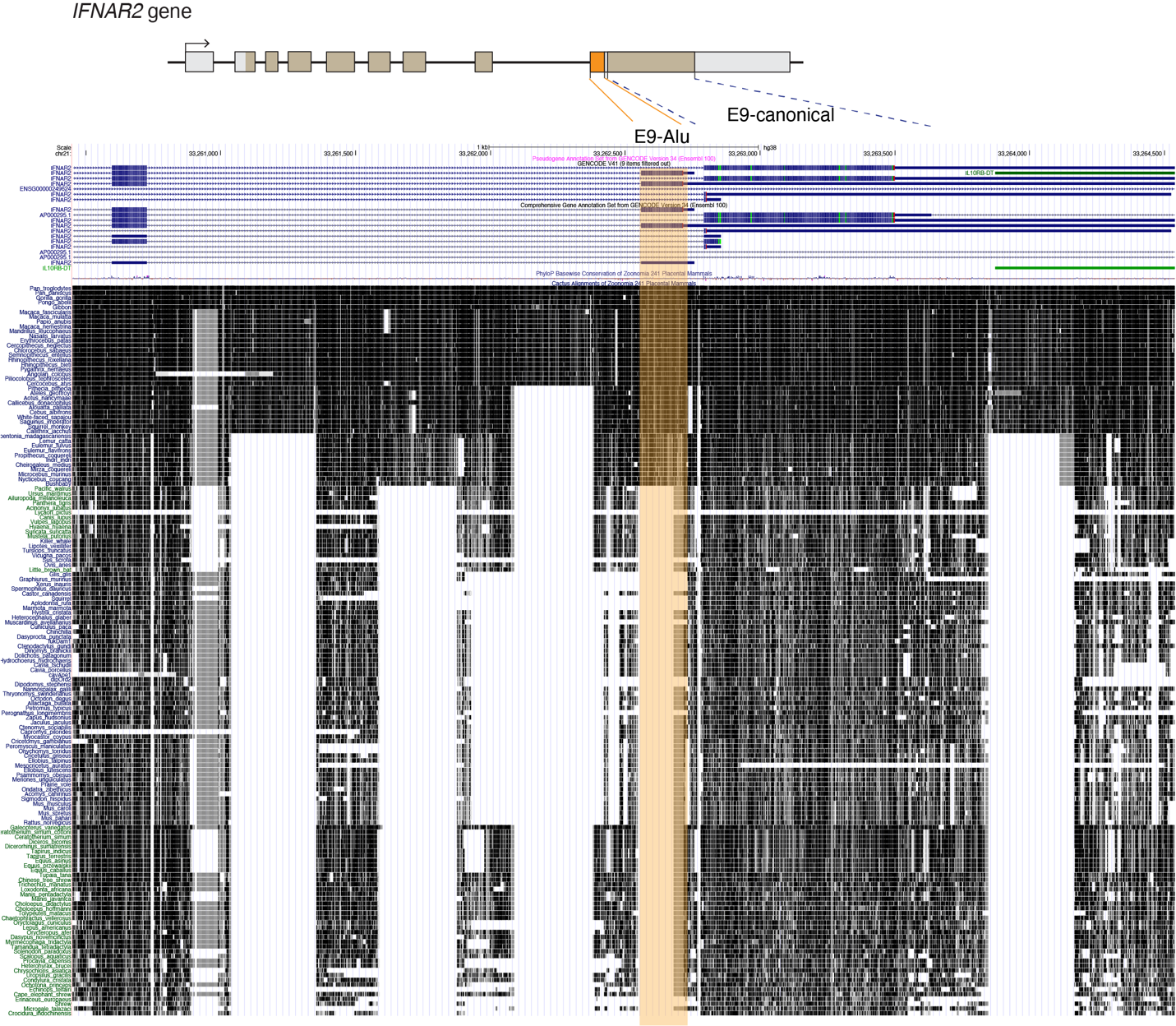

**Fig. S21) 241-way Vertebrate Genome Alignment at the *IFNAR2* locus.**

Screenshot of the human IFNAR2 locus (exon 8 to exon 9) superimposed to a whole-genome alignment of 241 vertebrate species (Zoonomia Consortium) from the UCSC genome browser. Highlighted in orange is the E9-Alu exon, conserved uniquely among simian primates. In prosimian monkeys the syntenic locus is occupied by an ancestral fossil Alu (FAM), which still retains sequence homology to the simian-specific Alu-Jr. The syntenic locus is not conserved in other vertebrate species.

Supplementary Figure 22

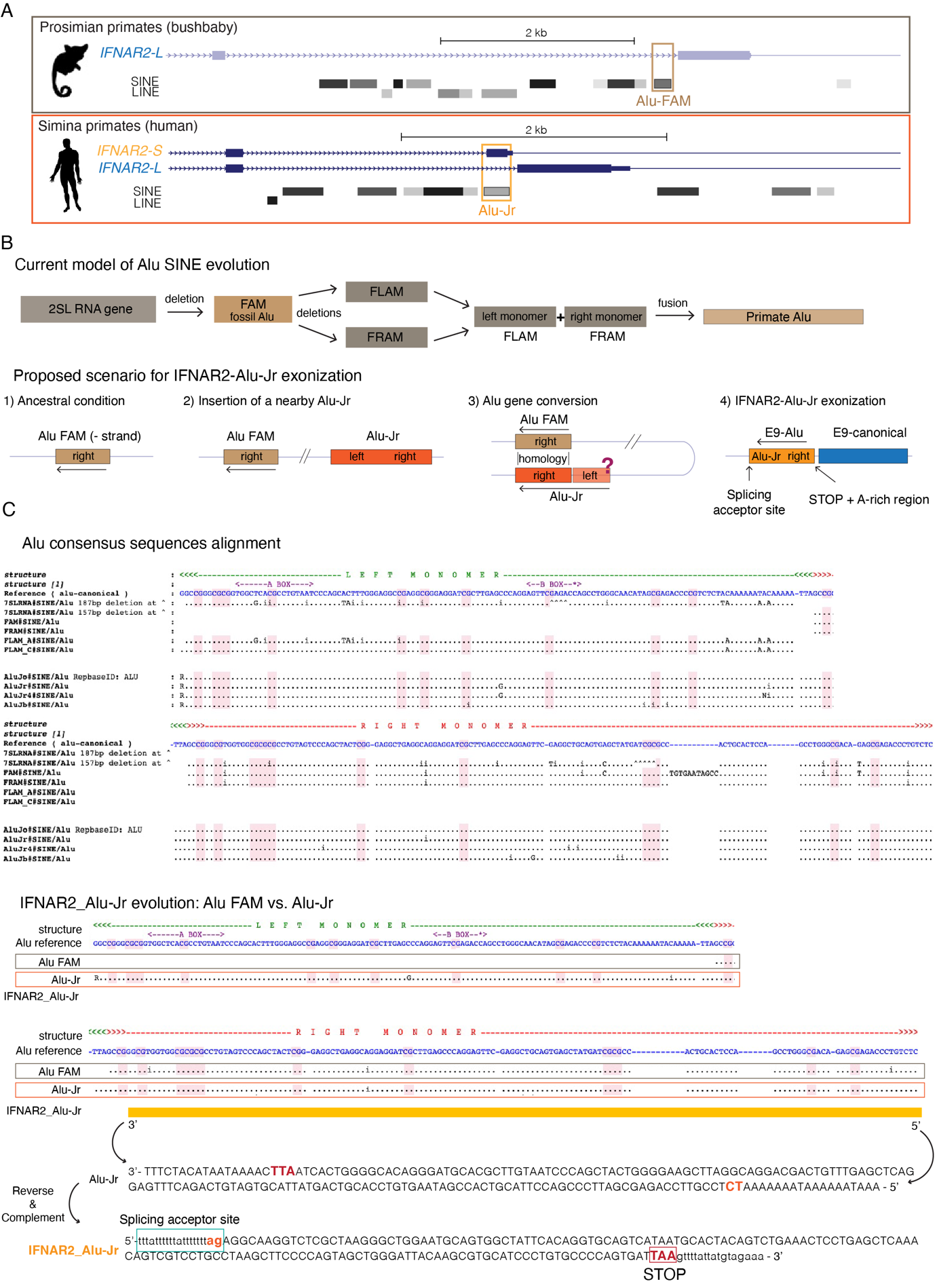

**Fig. S22) Proposed model of IFNAR2-S evolution in simian primates. A)** Screenshot of the IFNAR2 locus in a simian primate (human) and a prosimian primate (bushbaby) showing that the IFNAR2-S isoform is specific to simian primates. The exonized Alu-Jr present in simian primates replaced the synthenic ancestral Alu FAM present in prosimian primates through a gene conversion event. **B)** **Top**: Schematic model of Alu SINE evolution. Adapted from: 10.1093/nar/20.13.3397 and 10.1186/gb-1192 2011-12-12-236. **Bottom**: Proposed model of the Alu gene conversion event leading to the exonization of the IFNAR2_Alu-Jr in simian primates. **C)** Multiple sequence alignment of consensus Alu sequences was taken from:

http://www.repeatmasker.org/AluSubfamilies/humanAluSubfamilies.html. At the bottom the

IFNAR2_Alu-Jr sequence is provided in reference to the Alu consensus sequence alignment.

Supplementary Figure 23

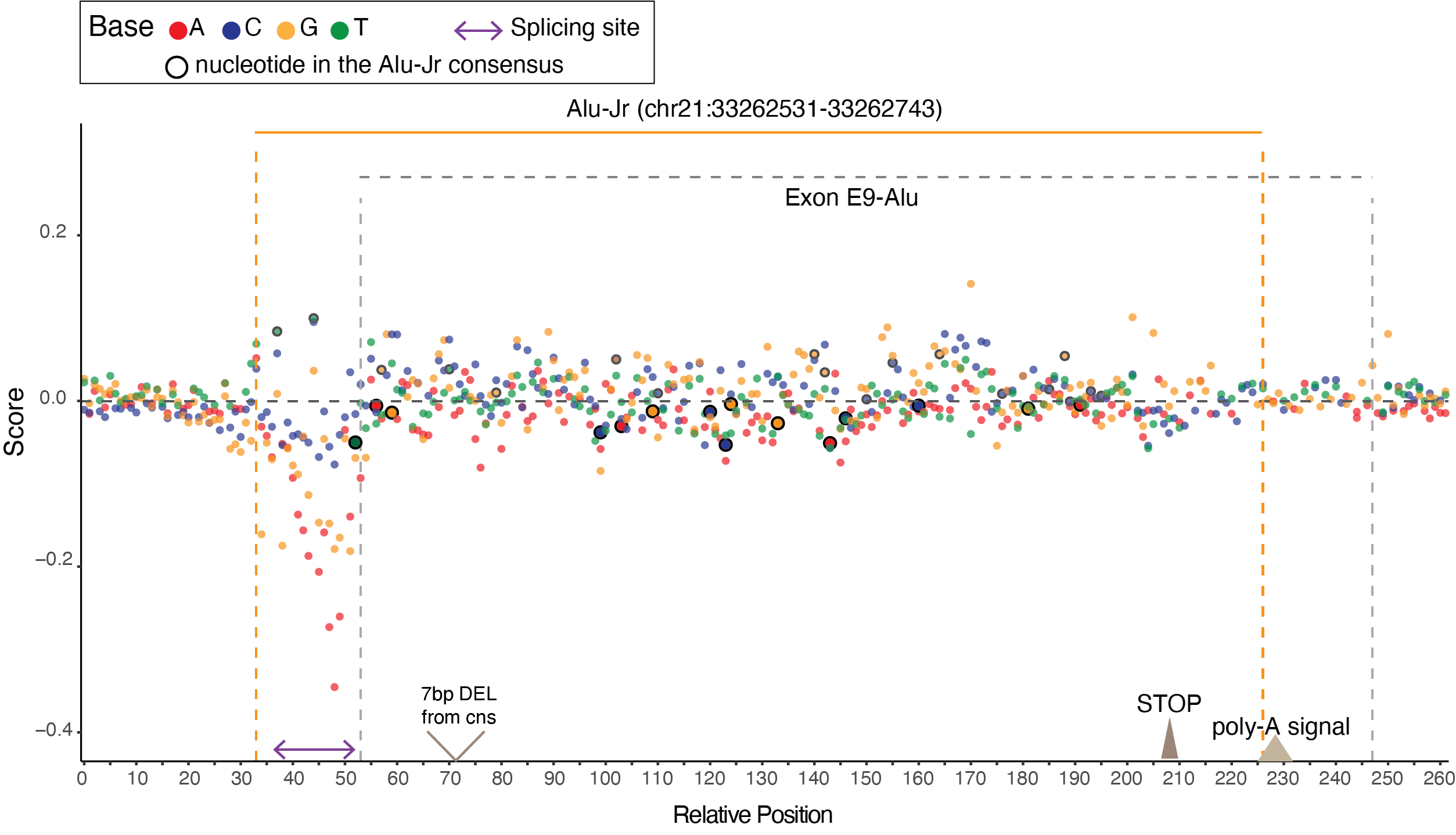

**Fig. S23) Mutation spectrum of the exonized Alu-Jr.** The software *Alubaster* (https://github.com/splicebox/Alubaster) was used to calculate the weight of single nucleotide mutations for each position of the Alu-Jr SINE (+/-35bp) on splicing. Nucleotide changes with a negative score are predicted to negatively affect splicing of the alternative exon E9-Alu, and are particularly concentrated at the splicing acceptor site region. Additionally, we aligned the sequence of the modern Alu-Jr to the Alu-Jr SINE consensus sequence to identify mutation that allowed and/or positively affected the alternative splicing of exon E9-Alu in simian primates. Compellingly, one such mutation (SNP ID: 21:33262573 G>A; relative position 66) is associated with severe COVID-19, and predicted to have a negative score.
